# MEK inhibition enhances presentation of targetable MHC-I tumor antigens in mutant melanomas

**DOI:** 10.1101/2022.01.10.475285

**Authors:** Lauren E. Stopfer, Nicholas J. Rettko, Owen Leddy, Joshua M. Mesfin, Eric Brown, Shannon Winski, Bryan Bryson, James A. Wells, Forest M. White

**Affiliations:** Department of Biological Engineering, Massachusetts Institute of Technology, Cambridge, MA 02139, USA; Koch Institute for Integrative Cancer Research, Center for Precision Cancer Medicine, Massachusetts Institute of Technology, Cambridge, MA 02139, USA; Department of Pharmaceutical Chemistry, University of California San Francisco, San Francisco, CA, 94158, USA; Ragon Institute of MGH, MIT, and Harvard, Cambridge, Massachusetts, 02139, USA; Pfizer Boulder Research & Development, Boulder, CO 8030, USA; Chan Zuckerberg Biohub, San Francisco, CA, 94158, USA

**Keywords:** MHC-I, Antigen presentation, immunopeptidomics, mass spectrometry, kinase inhibitor, melanoma

## Abstract

Combining multiple therapeutic strategies in NRAS/BRAF mutant melanoma – namely MEK/BRAF kinase inhibitors, immune checkpoint inhibitors, and targeted immunotherapies – may offer an improved survival benefit by overcoming limitations associated with any individual therapy. Still, optimal combination, order, and timing of administration remains under investigation. Here, we measure how MEK inhibition (MEKi) alters anti-tumor immunity by utilizing quantitative immunopeptidomics to profile changes in the peptide MHC (pMHC) repertoire. These data reveal a collection of tumor antigens whose presentation levels are selectively augmented following therapy, including several epitopes present at over 1000 copies-per-cell. We leveraged the tunable abundance of MEKi-modulated antigens by targeting 4 epitopes with pMHC-specific T cell engagers and antibody drug conjugates, enhancing cell killing in tumor cells following MEK inhibition. These results highlight drug treatment as a means to enhance immunotherapy efficacy by targeting specific upregulated pMHCs and provide a methodological framework for identifying, quantifying, and therapeutically targeting additional epitopes of interest.

**SIGNIFICANCE:** Kinase inhibitor treatment in NRAS/BRAF mutant melanoma can sensitize tumors to immunotherapy, in part through an increase in average surface presentation of peptide MHC molecules. Here, we demonstrate that MEK inhibition selectively boosts epitope abundance of select tumor-associated antigens *in vitro* and *in vivo*, enhancing targeted immunotherapy efficacy against these treatment-modulated epitopes.

## INTRODUCTION

In recent years, cancer treatment paradigms have increasingly incorporated information regarding a patient’s genetic profile to identify appropriate therapeutic modalities, otherwise known as “precision medicine.” Targeted therapies against aberrant activation of the mitogen-activated protein kinase (MAPK) signaling pathway, including BRAF and MEK inhibitors (BRAFi, MEKi), have transformed the standard of care for *BRAF* and *NRAS* mutant melanoma patients - representing ~50% and ~20% of melanomas, respectively (1, 2). Unfortunately, despite these targeted therapies showing some initial efficacy in extending progression free survival (PFS), either alone (MEKi, *NRAS*) or in combination (*BRAF*), a majority of patients acquire resistance and experience disease progression within one year (3–9). Immune checkpoint inhibitors (ICIs), which target cell surface receptors controlling the activation or inhibition of an immune response, have shown remarkable clinical success in melanoma (10, 11). However, only a subset of patients respond, and those who do frequently experience immune related adverse events (irAEs) and many develop resistance (12, 13).

It has been proposed that combining MAPK inhibitors and ICIs may increase efficacy, in part due to increasing evidence that MEK/BRAF inhibitors can sensitize tumors to immunotherapy through upregulation of class I major histocompatibility molecules (MHCs), as well as increase immune cell infiltration, T cell activation, antigen recognition, and more (14–17). NRAS-mutant melanoma trials have suggested that MEKi/ICI treatment may enhance PFS (9, 18). Additionally, several clinical trials evaluating a triple combination of MEKi, BRAFi, and ICIs have shown enhanced efficacy in BRAF-mutant melanoma, though at the expense of increased toxicity (19, 20). Therefore, despite promising initial results, there remains much to learn about how exposure to kinase inhibitors alters the immune system, and how these alterations can be leveraged with ICIs and/or targeted immunotherapies (21). Measuring how the antigen repertoire, referred to as the “immunopeptidome,” presented by class I MHCs changes in response to therapy is central to understanding the relationship between drug treatment and immune response, as recent reports highlight the potential for dynamic repertoire shifts in the identify and abundance of peptide MHCs (pMHCs) following perturbation (22–24). To better understand how to optimally combine therapies in BRAF/NRAS melanoma and identify antigens as therapeutic targets, a precise, molecular understanding of relative and absolute quantitative changes in pMHC presentation following treatment is required.

To this end, we used quantitative immunopeptidomics to measure the relative changes in presentation of pMHC repertoires in response to MEKi in vitro and in vivo. This analysis showed increased expression of both putative and well-characterized tumor associated antigens (TAA) following MEKi treatment. To interrogate the mechanisms underlying altered pMHC repertoires, we performed a quantitative multi-omics analysis and integrated the results with quantitative immunopeptidomic data to identify associations between intracellular response to MEKi and extracellular immune presentation. This analysis suggested the selective modulation of melanoma differentiation antigens and other TAAs by MEKi though a shared mechanism, highlighting potential antigen targets for targeted immunotherapy whose expression can be tuned with MEK inhibitor treatment.

Copies-per-cell estimations of 18 MEKi-modulated TAAs enabled the selection of four TAAs with high MEKi-induced expression as targets for pMHC-specific antibody-based therapies, which show enhanced ability to mediate T cell cytotoxicity with higher antigen expression levels (25–28). The pMHC-Abs were used to generate antibody-drug conjugates and T-cell engagers, which reveal a strong relationship between epitope density, therapeutic modality, and cytotoxicity, and highlight MEKi as a means to enhance efficacy by increasing target antigen expression. Our work provides the methodological framework to discover and exploit highly expressed drug-induced pMHC complexes for new immunotherapies.

## RESULTS

### MEK inhibition increases MHC-I expression in melanoma cell lines

To evaluate how MEK inhibition alters pMHC expression in *NRAS* and *BRAF* mutant melanomas, we selected 2 *NRAS* and 4 *BRAF* mutant cell lines (V600E) which exhibited a range of sensitivities to binimetinib (**Fig. S1A**). We measured class-I MHC (MHC-I) surface expression with flow cytometry and found 72 hours of treatment resulted in a maximal increase in expression over a DMSO treated control without requiring cell passaging (**Fig. S1B**). Hence, we selected 72 hours as the timepoint for all subsequent experiments. All cell lines showed elevated surface MHC-I expression following low (100 nM) or high-dose (1 *μ*M) binimetinib treatment at 72 hours, with high-dose treatment generally resulting in a larger increase (**Fig. 1A, Fig. S1C**). Primary melanocytes treated with binimetinib did not show a strong change in surface HLA expression, similar to previously reported results in trametinib-treated PBMCs (14), suggesting this effect is specific to oncogenic cell phenotypes with amplified MAPK signaling (**Fig. S1D**).

**Fig. 1.**
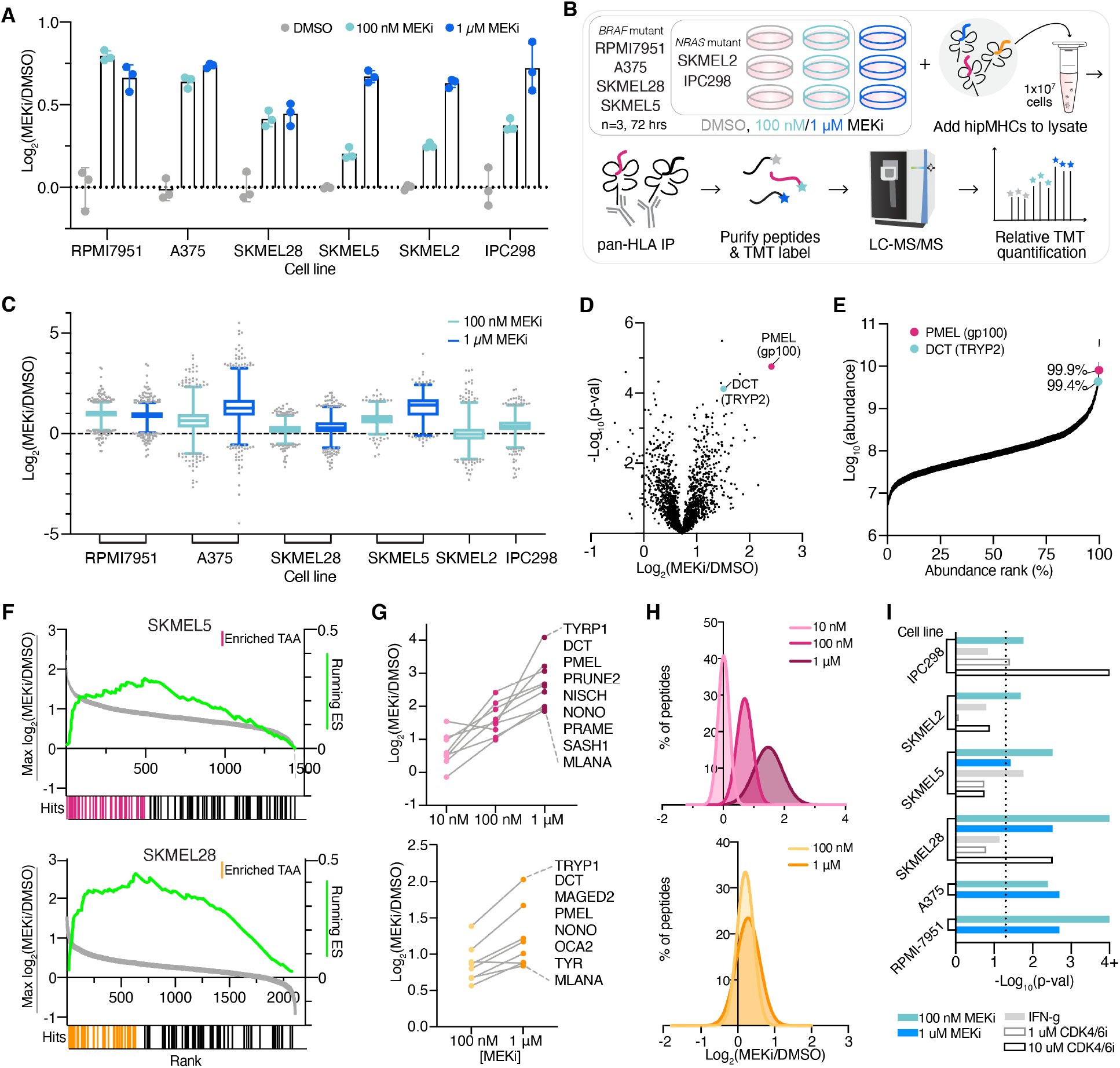
MEKi enriches TAA presentation on pMHCs. **(A)** Fold change in median surface expression levels (over average DMSO control condition) of HLA-A/B/C in cell lines treated with vehicle control or binimetinib (MEKi) for 72 hr. Error bars represent standard deviation of n=3 biological replicates. **(B)** Experimental setup for quantitative immunopeptidomics experiments. **(C)** Relative changes in pMHC expression +/- MEKi. Data are represented as a box and whiskers plot, with whiskers displaying the 1-99 percentiles. **(D)** Volcano plot of the average fold change in pMHC expression for SKMEL5 cells treated with 100 nM binimetinib for 72 hr (n=3 biological replicates for DMSO and MEKi treated cells) versus significance (mean-adjusted p value, unpaired two-sided t test). **(E)** pMHCs ranked by precursor ion area abundance. **(F)** TAA Enrichment plots of TAA enrichment in SKMEL5 +/- 100 nM MEKi (top, pink) and SKMEL28 +/- 100 nM MEKi (bottom, orange), displaying running enrichment scores (green, right y-axis), and fold change in pMHC presentation (left y-axis) versus rank (x-axis) for each peptide (gray). Hits denote TAA peptides, and colored hits represent enriched TAAs. SKMEL5 p = 0.001=4, SKMEL28 p = 0.001. **(G)** Selected enriched TAA peptides in SKMEL5 (top) and SKMEL28 (bottom) analyses. **(H)** Frequency distribution of pMHC fold change with MEK inhibition. SKMEL5 (top): 10 nM: μ=0.01, 100 nM: μ=0.70, 1 μM: μ=1.47. SKMEL28 (bottom): 100 nM: μ=0.21, 1μM: μ=0.28. **(I)** Significance values for TAA pathway enrichment. Dotted line indicates and p<0.05 and values ≥4 (Log_10_ adjusted) represent p<0.0001.

We next investigated how the pMHC repertoires presented on these six cell lines were altered quantitatively in response to MEKi treatment. We employed our previously described framework for multiplexed, quantitative profiling of pMHC repertoires utilizing isobaric labeling (TMT) and heavy isotope-labeled peptide MHCs (hipMHC) standards for accurate relative quantitation of endogenous pMHCs. In triplicate, cells were treated with DMSO or binimetinib (100 nM *NRAS* mutant cells, 100 nM/1 *μ*M *BRAF* mutant cells) for 72 hours (**Fig. 1B**). Cells were lysed, and three hipMHC standards were spiked into the lysate mixture prior to immunoprecipitation. Isolated endogenous and isotopically labeled peptides were subsequently labeled with TMT, combined, and analyzed by LC-MS/MS for quantitative immunopeptidomic profiling (**Dataset S1**). Peptides matched expected class I length distributions (**Fig. S2A)**, and a majority were predicted to be binders of each cell line’s HLA allelic profile (**Fig. S2B, SI Appendix, Table S1**).

Quantitative immunopeptidomics showed a median increase in pMHC expression levels following binimetinib treatment in most conditions, with similar average changes observed across peptides predicted to bind to HLA-A, HLA-B, and HLA-C (**Fig. S2C**). However, in contrast to-surface staining, measuring the average change in HLA expression, MS analysis showcased a wide distribution in presentation levels across peptides (**Fig. 1C, Fig. S3**). For example, while the average fold change in HLA levels in A375 cells treated with 1 *μ*M binimetinib was 2.45-fold, some peptides increased 16-fold or more in presentation while others decreased 4-fold. In SKMEL2 cells, several peptides changed 3 to 4-fold in presentation despite no change in surface HLA expression. These data illustrate the highly dynamic nature of the immunopeptidome, where individual pMHCs experience significant changes in presentation often not captured by surface staining alone.

### Tumor associated antigens are selectively enriched in presentation with MEK inhibition

We investigated which peptides increased significantly relative to the median change in presentation to determine if any pMHCs were selectively enriched following MEK inhibition. We observed that two peptides in the SKMEL5 (low-dose MEKi) analysis, derived from known TAAs (dopachrome tautomerase [DCT or “TYRP2”] and premelanosome protein [PMEL or “gp100”]), had high changes in presentation, increasing 2.8 and 5.3-fold, respectively (**Fig. 1D**). These peptides were also highly abundant, ranking in the 99^th^ percentile of precursor ion abundance (**Fig. 1E**).

To evaluate whether enriched presentation of DCT and PMEL peptides was indicative of increased expression of TAA-derived peptides following MEKi treatment broadly, we performed a non-parametric test to measure TAA enrichment significance. For this analysis, we compiled a custom TAA library derived from the literature and online databases, (**SI Appendix, Table S2**) (29-32) utilizing the peptide’s source proteins to generate a protein-based TAA library to accommodate peptides derived from all alleles (**SI Appendix, Table S3**). Peptide source proteins were rank-ordered by fold change in presentation with MEKi; in cases where multiple peptides were derived from the same source protein, the maximal/minimal fold change was selected to assess positive/negative enrichment.

In both SKMEL5 and SKMEL28 cells, TAAs were significantly positively enriched following low-dose MEKi (**Fig. 1F**). Beyond DCT, and PMEL, enriched TAAs included melanoma differentiation antigens from the MAGE family, MLANA (MART-1), and TYR, representing well characterized antigens with demonstrated immunogenic potential (**Fig. S4A**) (33, 34). Many TAAs showed a dose dependent increase in presentation, occurring regardless of whether mean HLA expression increases proportionally (**Fig. 1G-H).** Even sub-cytotoxic doses of MEKi (10 nM) resulted in an increase in TAA presentation despite no change in average MHC surface expression (**Fig. S4B**).

This effect was not exclusive to binimetinib, as trametinib-treated SKMEL5 cells showed similar TAA enrichment (**Fig. S4C**). Peptides rank-ordered by precursor ion abundance also reached significance, suggesting TAAs are both some of the most abundant peptides presented (Fig. S4D). Applying the enrichment analysis framework to all cell lines and treatments revealed MEKi treatment significantly enriched (p<0.05) TAA presentation in all cases, suggesting a mechanistic basis for this response (**Fig. 1I**). Other perturbations such as IFN-γ stimulation and CDK4/6 inhibitor treatment have also been shown to increase antigen presentation, however the TAA enrichment analysis applied to these previously published datasets revealed only a minority of cell lines/treatment conditions showed significant TAA enrichment **(Fig. 1I)** (24). Taken together, these data suggest MEK inhibition causes a distinct peptide repertoire shift from IFN-γ stimulation, robustly driving TAA upregulation distinct from other perturbations.

### Cell line xenografts show enhanced TAA presentation following MEK and BRAF inhibition *in vivo*

We next evaluated whether TAA enrichment following MEK inhibition translated *in vivo* at early timepoints. Four melanoma cell lines were inoculated subcutaneously in immunocompromised mice, and mice were treated with vehicle control or binimetinib for 1, 2, 3, or 5 days in triplicate prior to tumor harvesting (**Fig. 2A**). For *BRAF* mutant lines, three additional mice were treated for 3 days with encorafinib (BRAF inhibitor, BRAFi) or encorafinib and binimetinib as a combination therapy. Class-I pMHCs from tumors were isolated and subsequently profiled by quantitative multiplexed immunopeptidomics (**Dataset S2, Fig. S5A-B**).

**Fig. 2.**
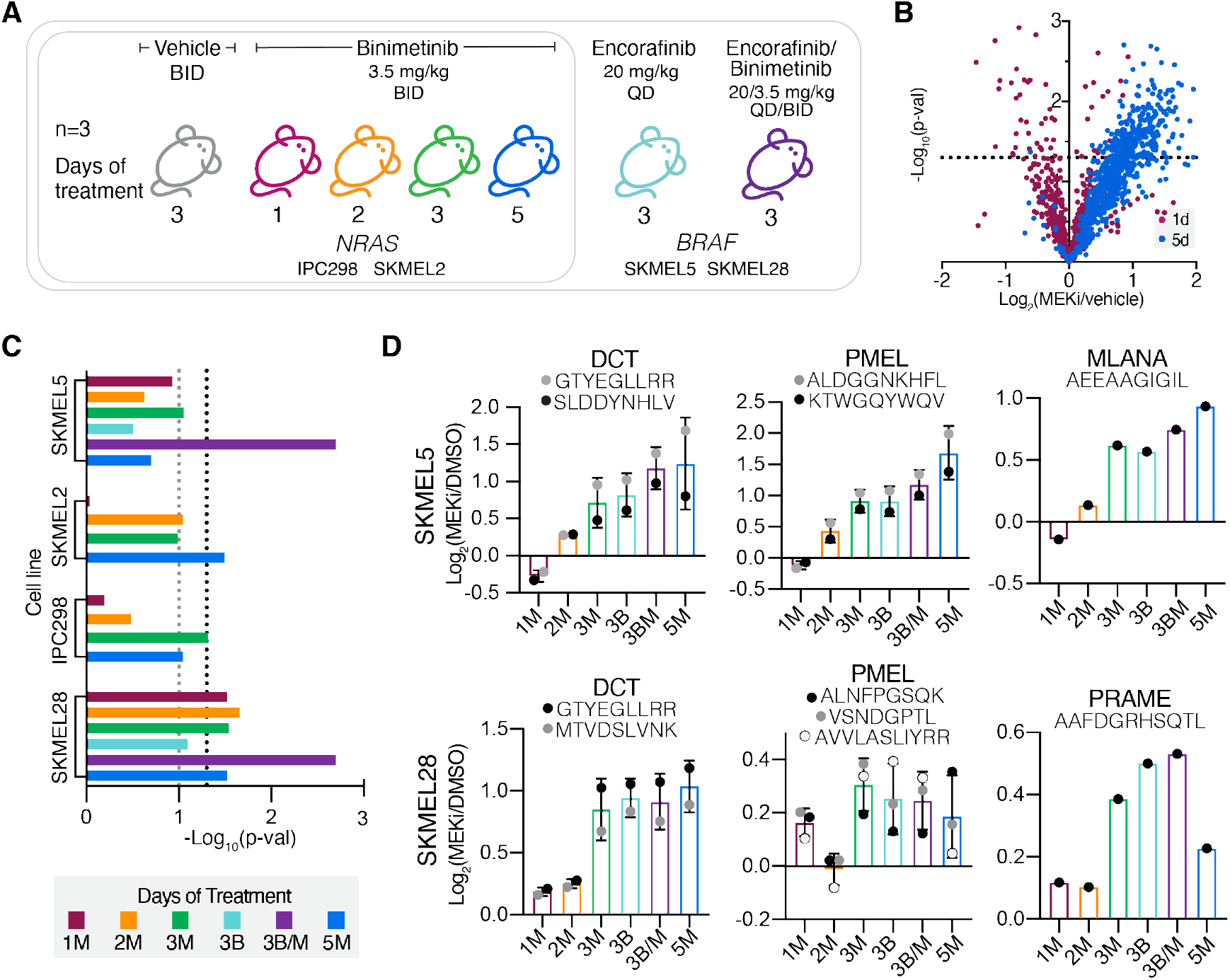
In vivo analysis of TAA pMHC enrichment in cell line xenografts. **(A)** Experimental setup for cell line xenograft studies of mice + binimetinib or encorafinib. X-axis describes days of therapy and drug treatment (M=MEKi, binimetinib and B=BRAFi, encorafinib). **(B)** Volcano plot, average fold change in pMHC expression with binimetinib treatment (n=3 biological replicates for DMSO and MEKi treated cells) versus significance (mean-adjusted p value, unpaired two-sided t test) for IPC298 CLX. **(C)** TAA enrichment significance values for each analysis. Black dotted line represents p≥0.05, grey = p≥0.01. **(D)** Changes in pMHC expression for select melanoma differentiation antigens. Errors bars represent standard deviation when >1 peptide from each source protein was identified.

Among cell line xenografts (CLXs), treatment with binimetinib for just 1 or 2 days minimally altered mean HLA presentation levels, with maximal changes in presentation observed after 3 or 5 days of treatment (**Fig. 2B, Fig. S5C**). In *BRAF* mutant CLXs, combination therapy showed higher (SKMEL5) or similar (SKMEL28) changes in pMHC presentation compared to MEKi monotherapy, suggesting combination therapy in BRAF tumors may further improve antigenicity of tumors in some cases.

We next performed TAA enrichment analysis and observed significant enrichment in at least one treatment condition across CLXs, with SKMEL28 CLXs showing robust enrichment across timepoints (**Fig. 2C**). Melanoma differentiation antigens showed positive increases in presentation following MEK and BRAF inhibition across all cell lines, often above median foldchanges (**Fig. 2D, Fig. S5D**). For example, the PMEL peptide “ALDGGNKHFL” had a nearly four-fold increase in presentation after 5 days of MEKi in SKMEL5 CLXs, far exceeding the median pMHC fold-change value of 1.15.

Finally, we performed TAA enrichment analysis with peptides rank ordered by peak area abundance for SKMEL5 and SKMEL28 CLX samples and found that both showed significant enrichment (p<0.0001), as seen in the *in vitro* analyses (**Fig. S5E**). High abundance TAAs mapped to peptides that had some of the highest changes in expression following MEKi (**Fig. S5F**), further confirming our observations that TAAs are some the most abundant and differentially expressed pMHCs following MEK inhibition.

### EMT-TF switching drives MITF and melanoma differentiation antigen expression

To assess whether MEKi-induced, enriched TAA pMHC presentation could be predicted using other datatypes, we performed a multi-omics analysis and compared changes in protein and transcript expression, as well as changes in ubiquitination as a proxy for protein degradation, following MEKi treatment to changes in pMHC presentation (**Fig. S6A**, **Dataset S3-5**). Overall, there was no significant correlation between changes in pMHC expression and transcript/protein/ubiquitination after MEK inhibition, suggesting these datatypes cannot necessarily be used exclusively for predicting pMHC repertoire alterations (**Fig. 3A, Fig. S6B**). For example, vimentin has increased pMHC presentation but decreased transcript and protein expression, suggesting the elevated pMHC expression is likely due to other post-translational processing such as enhanced degradation as measured by ubiquitination (**Fig. 3B**).

**Fig. 3.**
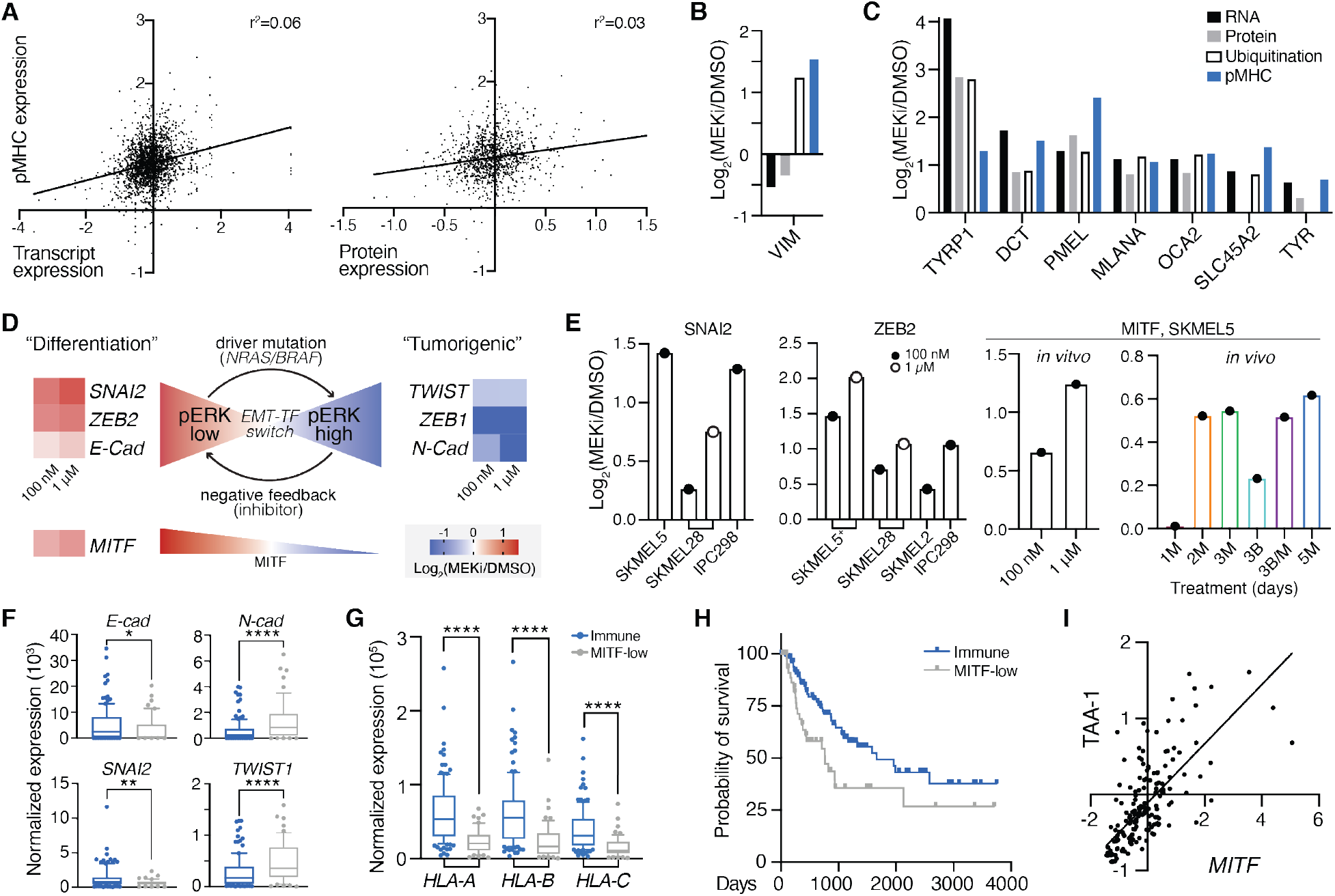
MEKi-induced EMT phenotype switching enhances TAA expression. **(A)** Correlation between pMHC expression and transcript expression (left) / protein expression (right) in SKMEL5 cells +/- 100 nM MEKi. Values represent Log_2_(MEKi/DMSO). **(B)-(C)** Changes in RNA, protein, ubiquitination, and pMHC expression of selected targets. **(D)** Schematic of EMT-TF switching, with changes in transcription (SKMEL5 +/100 nM or 1 uM binimetinib) shown next to gene names. **(E)** Maximum change in expression of pMHCs derived from EMT-related source proteins. M = MEKi, B = BRAFi. **(F)-(G)** Normalized expression of select genes in NRAS/BRAF mutant immune (blue, n=122) and MITF-low (grey, n=54) classified tumors. *p<0.05, **p<0.01, ****p<.0001, unpaired 2-tailed t-test. **(H)** Kaplan Meier curve of BRAF/NRAS immune and MITF-low melanomas. p=0.012, log-rank test. **(I)** Correlation in expression of MITF and average TAA-1 gene set Z-scored expression for immune/MITF-low tumors, r=0.723, p<0.0001 (two-tail).

Despite the lack of general correlation, a clustering analysis of changes in pMHC, protein, and RNA expression following MEKi revealed a subset of source genes including DCT, PMEL, TYR, TRYP1, SLC45A2, and others, which showed increases across all datatypes (including ubiquitination when available) (**Fig. 3C**, **Fig. S6C-D**). This result demonstrates a clear connection between pMHC presentation and changes in transcription, translation, and degradation in the context of a subset of melanoma differentiation antigens.

To investigate the underlying biological mechanism responsible for select TAA enrichment, we performed gene set enrichment analysis (GSEA) against the cancer hallmarks pathway database and found significant negative enrichment of the epithelial to mesenchymal transition (EMT) pathway (**Fig. S6E**) (35).

Transcript expression patterns in SKMEL5 cells provide evidence for a previously described EMT program in melanoma termed “EMT-transcription factor (TF) switching,” where SNAI2 and ZEB2 act as oncosuppressive proteins during melanocyte differentiation under MITF control (36, 37). In response to MAPK pathway activation, EMT-TFs ZEB1 and TWIST are upregulated to promote dedifferentiation and tumorigenesis, and inhibition of this pathway (i.e. MEK or BRAF inhibitors, low phospho-ERK) can reverse the EMT-TF phenotype back to a “differentiation” state (high MITF). Here, cells showed increased “differentiation” and decreased “tumorigenic” marker transcript expression following MEK inhibition (**Fig. 3D**).

Previous studies have demonstrated that quantitative changes in pMHC repertoires reflect biological response to perturbation (24, 38). In line with this finding, we find pMHCs derived from differentiation-associated proteins like SNAI2, ZEB2, and MITF increasing in presentation following MEKi in *in vitro* and *in vivo* analyses, further connecting the intracellular response to treatment to extracellular immune presentation (**Fig. 3E, Fig. S7**).

We next queried TCGA transcriptional data of 176 BRAF or NRAS mutant cutaneous melanoma patients to evaluate whether there was a relationship between melanoma differentiation antigen expression and EMT-TF phenotypes (39). Tumors were previously classified into subclasses by Akbani et al., including “immune” for tumors with high immune infiltration and “MITF-low” for low MITF and target gene expression, whose EMT-TF expression profiles matched the previously reported phenotypes (**Fig. 3F**) (36). MITF-low tumors had significantly lower HLA-A/B/C expression and a lower probability of survival, and MITF expression was significantly correlated with melanoma differentiation antigen expression (**Fig. 3G-I**). These data suggest “MITF-low” BRAF/NRAS tumors may benefit from MEKi or other MAPK pathway inhibitors to induce a high-MITF, “differentiation” phenotype and suggest a common mechanism to augment TAA pMHC expression in melanoma.

### Absolute quantification of treatment-modulated tumor associated antigens

MEK inhibitor-modulated TAAs present an attractive class of epitopes for targeted immunotherapy, as these antigens have high abundance relative to other epitopes and their expression can be further augmented in response to therapy. We hypothesized MEKi may enhance the anti-tumor immune response for immunotherapies targeted against MEKi-modulated antigens, although determining the appropriate immunotherapeutic strategy for each antigen requires knowledge of epitope abundance (27, 28).

To this end, we performed absolute quantification experiments to estimate copies-per-cell abundance of 18 HLA-A*02:01 epitope targets that increase in presentation following MEKi in SKMEL5 cells. We utilized a previously developed assay, “SureQuant Iso-MHC” (40), where a series of three peptide isotopologues with 1,2, or 3 stable isotopically labeled (SIL) amino acids (1-3H) per target were loaded into MHC molecules (hipMHCs) and titrated into cell lysates across a 100-fold linear range as an embedded standard curve (**Fig. 4A**). A fourth isotopologue with 4 SIL-amino acids was added exogenously to leverage internal-standard triggered parallel reaction monitoring data acquisition (IS-PRM, “SureQuant”) for sensitive and selective targeting of endogenous peptides and embedded peptide standards. We estimated copies-per-cell for our 18 TAA panel in A375 and RPMI-7951 cells treated with DMSO, low, or high dose MEKi for 72 hours, and extended our previous analysis of SKMEL5 cells by measuring the target panel in SKMEL5 cells with high dose MEKi treatment and compared the data to DMSO and low dose measurements (40).

**Fig. 4.**
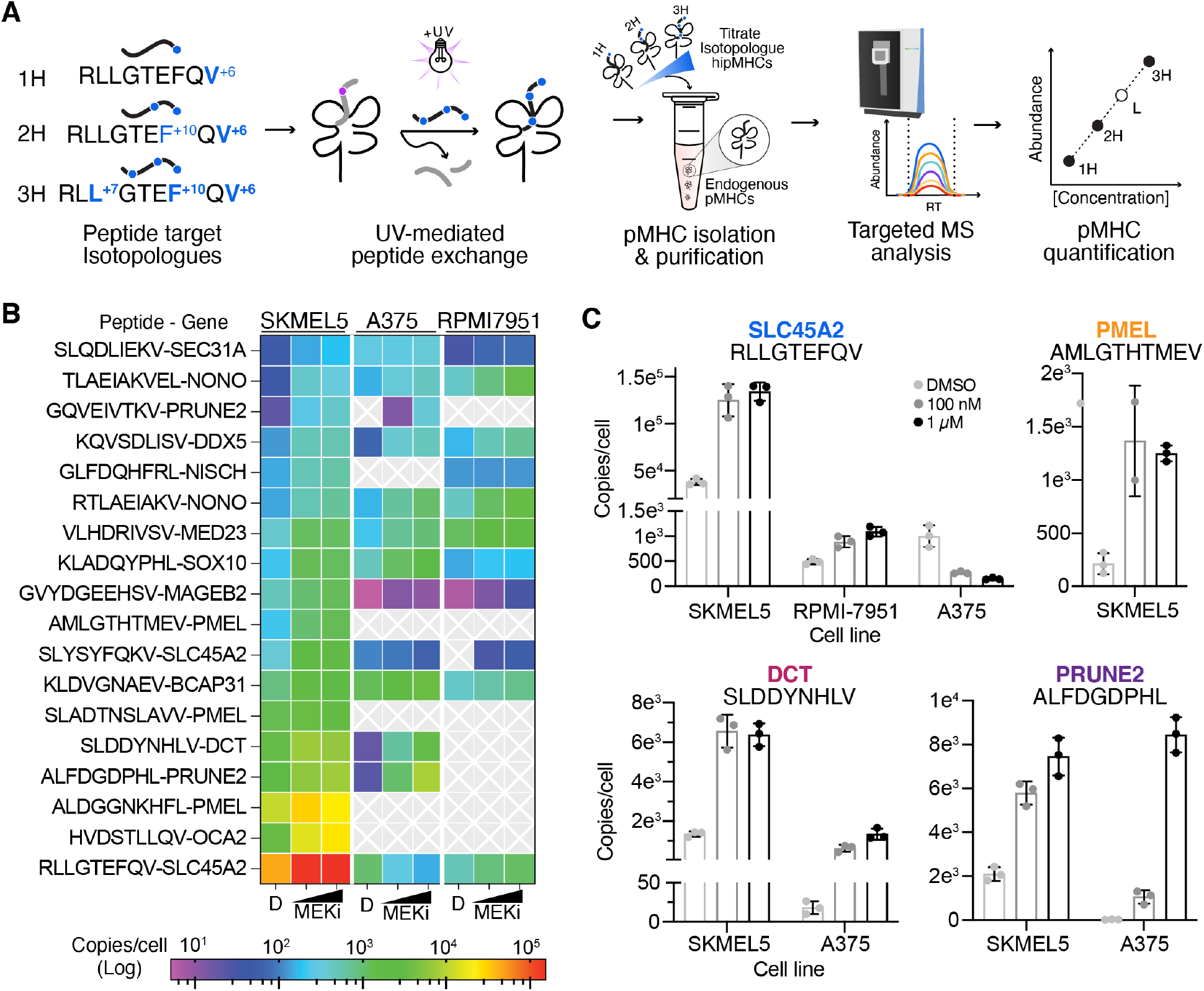
Absolute quantification of MEKi-inducible TAAs. **(A)** Schematic of SureQuant isoMHC workflow for multipoint absolute quantification of 18 TAAs. **(B)** Heatmap of copies/cell for each cell line treated with DMSO, 100 nM MEKi, or 1 μM MEK for n=3 biological replicates. **(C)** Copies/cell for select epitopes across cell lines and treatment conditions.

All 18 peptides were quantifiable across high MEKi treated SKMEL5 cells, as expected since the panel was developed using pMHCs identified in SKMEL5 cells (**Dataset S6**). While we would not expect to detect the entire panel across A375 and RPMI-7951 cell lines (for example, A375 cells are PMEL-), 13 and 11 peptides were quantifiable within A375 and RPMI-7951 cells, respectively (**Fig. 4B).** Copies-per-cell estimations spanned 5 orders of magnitude across peptides, cell lines, and treatments, highlighting the wide range in epitope abundances presented by cells.

### Generating pMHC-specific antibodies against MEKi-modulated TAAs

Antibody-based immunotherapies have shown the increasing promise of pMHC’s as therapeutic targets, both in the context of melanoma and cancer as a whole (25, 33, 41-43). MEKi-induction of shared TAAs described here may present a therapeutic opportunity to use pMHC-targeted antibodies in combination with MEKi. We selected four HLA-A*02:01 associated TAAs with high epitope abundance in SKMEL5 cells as antigens for antibody generation. These peptides (derived from SLC45A2, PMEL, DCT, and PRUNE2) exhibited a range of basal and MEKi-induced presentation levels - three of which were also identified in at least one other cell line (**Fig. 4C**). To identify pMHC-specific antibodies, we performed a phage display campaign first clearing 2 Fab-phage libraries with an immobilized pMHC containing a decoy peptide (GILGFVFTL from influenza, “Flu peptide”). Remaining phage were incubated with pMHC’s of interest and bound phage were eluted via TEV protease and subsequently propagated to enrich for selective binders (**Fig. 5A**). After iterative rounds of selection, ELISA screening of individual clones identified 15 unique Fabs that showed good specificity and high affinity (<20 nM) across our 4 pMHC targets (**Fig. S8**). Flow cytometry using T2 lymphoblasts – an HLA-A*02:01^+^ cell line null for TAP which allows for exogenous peptide loading – revealed 1 Fab per pMHC that specifically recognized the pMHC on the surface of cells in a peptide-dependent manner (**Fig. S9**). Upon conversion to IgG’s, these antibodies demonstrated exquisite selectivity in recognizing only peptide-specific target cells (**Fig. 5B**), each with subnanomolar affinity (**Fig. 5C**).

**Fig. 5.**
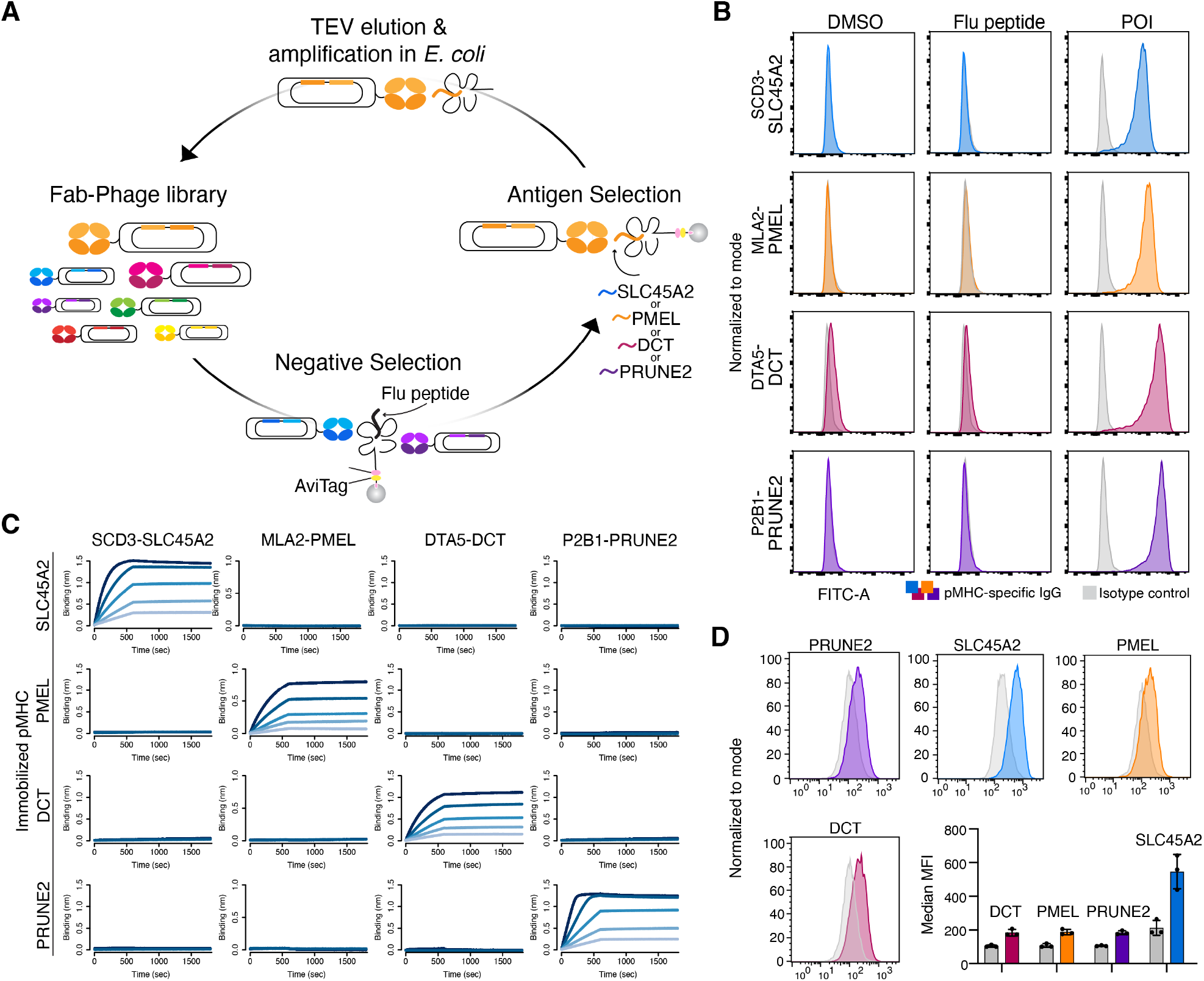
Generation of pMHC-specific antibodies. **(A)** Schematic of phage display selection. **(B)** Fluorescence intensity of T2 cells loaded with DMSO (negative control), a decoy FLU peptide, or peptide of interest (POI) stained with a pMHC-specific IgG (color) or an isotype control (grey). **(C)** Bio-layer interferometry (BLI) analysis of IgG’s (two-fold dilutions starting at 20 nM) against selected HLA-A*02:01 MHC-peptide complexes. **(D)** Fluorescence intensity of SKMEL5 cells treated with DMSO or 1 μM MEKi for 72 hours stained with Alexa fluor 488-conjugated pMHC-specific antibodies. Data show a representative histogram and bar graph of median MFI where error bars show standard deviation for n=3 biological replicates per condition.

SKMEL5 cells treated with DMSO or high dose MEKi for 72 hours displayed an increase in median fluorescence intensity in MEKi treated cell compared to DMSO when stained with fluorophore-conjugated pMHC-specific IgG’s, in line with our immunopeptidomic analysis. (**Fig. 5D**). Due to the superior tumor-specific expression profiles in skin (**Fig. S10**), as well poor biophysical properties of the antibody targeting the PRUNE2 pMHC (data not shown), we selected SLC45A2, DCT, and PMEL-specific antibodies to evaluate for efficacy *in vitro*.

### Therapeutic modality, antibody properties, and epitope expression influence efficacy of pMHC-specific antibody-based therapies

Previously reported data have demonstrated that ADCs targeting pMHCs require a high epitope density for efficacy, as only cells with expression levels generally above ~40,000 copies/cell showed an effect on viability greater than 20% (27). Here we hypothesized the high endogenous expression of the SLC45A2 “RLLGTEFQV” epitope in SKMEL5 cells may be effectively targeted by an ADC. To that end, we conjugated monomethyl auristatin F (MMAF), a tubulin polymerization inhibitor, to the anti-SLC45A2 pMHC IgG (**Fig. 6A**) and evaluated viability in SKMEL5 & RPMI-7951 (low epitope density) cells pre-treated with DMSO or 1 μM MEKi for 72 hours to augment pMHC presentation of the target epitope. In SKMEL5 cells, MEKi pretreatment resulted in a superior therapeutic window following 72 hours of ADC treatment, with a 40% reduction in viability achieved with MEKi compared to 28% with DMSO at 30 nM ADC (**Fig. 6B, Fig. S11A**). In contrast, RPMI-7951 cells showed just an 18% reduction in viability in both conditions (<1500 copies-per-cell), confirming that high epitope density is required for anti-pMHC ADC efficacy.

**Fig. 6.**
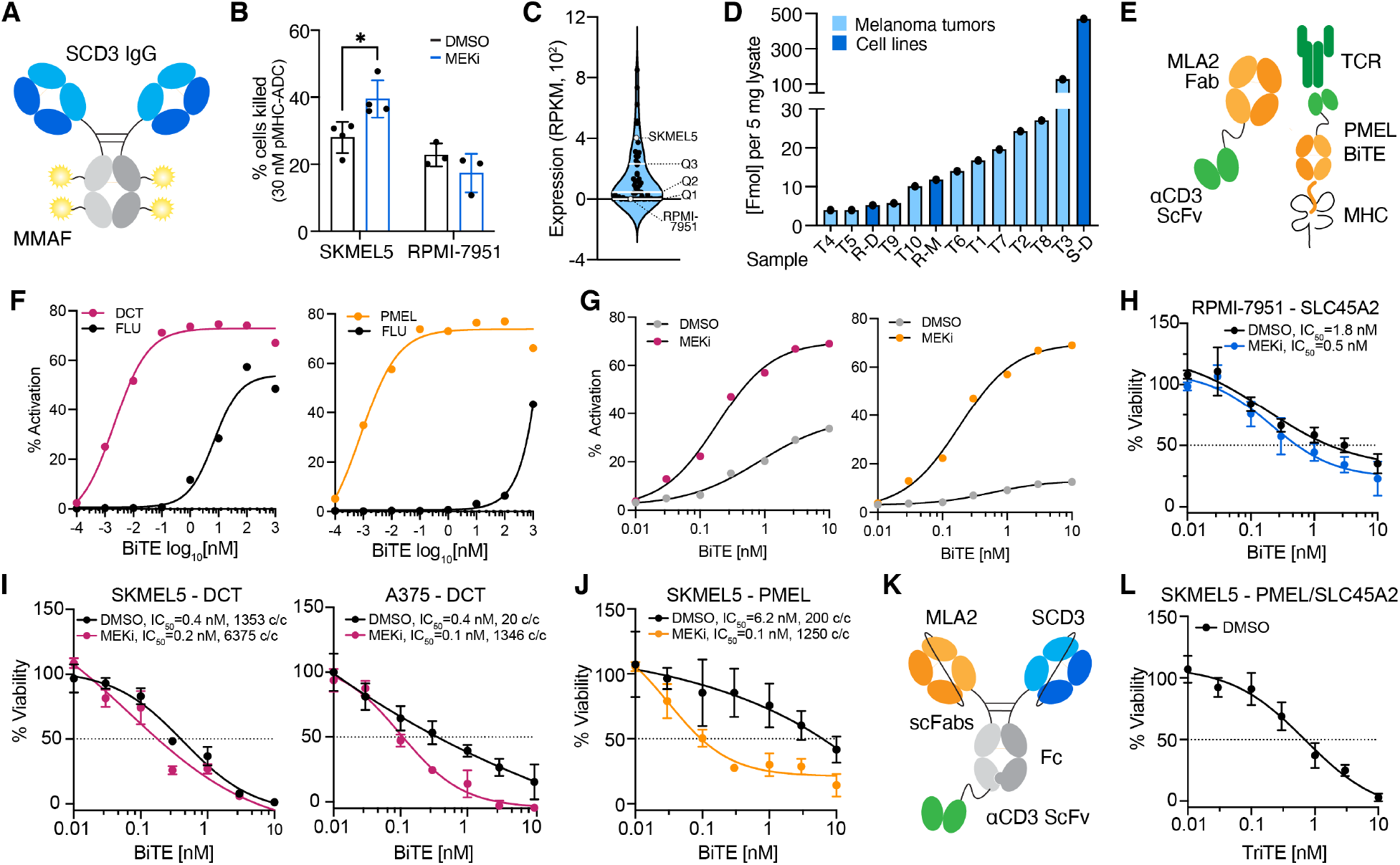
MEKi enhances cytotoxicity of pMHC-specific antibody-based therapies. **(A)** Schematic of ADC targeting the SLC45A2 epitope. **(B)** Percent of cells killed with SLC45A2-ADC relative to the DMSO control. Error bars represent +/- stdev for n=4 (SKMEL5) and n=3 (RPMI-7951) biological replicates. *adjusted p-value = 0.017, Sidak’s multiple comparisons test. **(C)** SLC45A2 transcript expression across n-57 BRAF/NRAS mutant melanoma cell lines. **(D)** RLLGTEFQV pMHC concentration across cell lines and melanoma tumors (previously reported). S=SKMEL5, R=RPMI-7951, D=DMSO, M=1 μM MEKi. **(E)** Schematic of PMEL-targeted BiTE. **(F)-(G)** Percent of GFP+ Jurkat cells following incubation with peptide-pulsed T2 cells (F) or SKMEL5 cells (G) and PMEL-BiTE. Lines represent a 4PL nonlinear fit, and error bars show +/- SEM for n=3 biological replicates. **(H)-(J), (L)** Cell viability (percentage of untreated control) of target cells incubated with normal human T cells (effector:target 2:1) & a pMHC-specific BiTE/TriTE for 48 hours. **(K)** Schematic of PMEL/SLC45A2-targeted TriTE.

Comparing SLC45A2 transcript expression across 57 BRAF/NRAS melanoma cell lines revealed SKMEL5’s expression is in the upper quartile of abundances, and RLLGTEFQV epitope concentration in SKMEL5 cells relative to previously profiled melanoma tumors (40) suggests the high SKMEL5 epitope abundance is unlikely to translate to a majority of patients (**Fig. 6C-D**). Therefore, while a subset of patients may benefit modestly from an ADC approach, an alternative strategy with a lower threshold for presentation may be more efficacious. PMEL and DCT epitopes showed lower surface presentation levels compared to SLC45A2 in SKMEL5 cells, thus we hypothesized bispecific T cell engagers (BiTEs) may be more potent against these epitopes, particularly in combination with MEKi (28, 44).

To this end, we generated BiTEs by fusing the PMEL, DCT, and SLC45A2 Fabs to the anti-CD3 single-chain variable fragment OKT3 (scFv, **Fig. 6E**). BiTE constructs showed selective T-cell activation in a NFAT-GFP Jurkat reporter cell line when incubated with T2 lymphoblasts loaded with target peptide in comparison to the decoy Flu peptide (**Fig. 6F, Fig. S11B**). We next tested Jurkat activation against SKMEL5 cells, and saw that cells pre-treated with 1 *μ*M MEKi for 72 hours showed superior activation across all 3 BiTES, suggesting higher target expression leads to a higher proportion of activated effector cells (**Fig. 6G, Fig. S11C**).

To assess cytotoxicity, we cocultured SKMEL5, RPMI-7951, or A375 cells pre-treated with DMSO or high-dose MEKi for 72 hours with primary human T cells isolated from healthy donor blood (effector to target ratio 2:1) in the presence of increasing concentrations of BiTE for 48 hours. While RPMI-7951 cells were not responsive to the SLC45A2-ADC, the SLC45A2-BiTE did yield a cytotoxic response, with MEKi pre-treated cells showing increased cell death with an IC_50_ of 0.5 nM with MEKi compared to 1.8 nM with DMSO (**Fig. 6H, Fig. S11D**). By comparison, SKMEL5 cells showed a similar response regardless of MEKi treatment, likely due to the already high presentation levels at baseline (**Fig. S11E**).

SKMEL5 cells showed a similar cytotoxic response to the DCT-BiTE regardless of treatment condition, possibly because both DMSO and MEKi-treated cells presented the target epitope at levels above 1000 copies/cell (**Fig. 6I**). For A375 cells, the DCT-BiTE showed a superior reduction in cell viability with MEKi-pretreatment, where expression levels increased from 20 to 1346 copies/cell. The PMEL-BiTE exhibited a similar trend in SKMEL5 cells, where a concentration of just 0.1 nM PMEL-BiTE was required to reduce SKMEL5 viability by 50% in MEKi pre-treated cells, in contrast to 6.2 nM required in DMSO-treated cells (**Fig. 6J**). These data suggest that epitopes presented above ~1000 copies/cell are most effectively targeted by BiTEs, and that MEKi treatment can be used to augment presentation levels for increased efficacy when endogenous expression of the target epitope is low.

In cases where MEKi treatment may not be a viable strategy to augment presentation of target antigens (ex. therapeutic resistance, non BRAF/NRAS mutant melanoma), utilizing a combination of BiTES that target patient epitopes may enhance cytotoxicity. Furthermore, two pMHC-specific antibodies can be combined to generate tri-specific T-cell engager molecules (TriTEs), which may increase cytotoxic response and/or lower the concentration of therapy required for efficacy. Accordingly, we generated a TriTE against SLC45A2 and PMEL (**Fig. 6K**) and observed enhanced Jurkat activation when T2 lymphoblasts were pulsed with both peptides at BiTE concentrations below 1 nM (**Fig. S11F**). In SKMEL5 cells (DMSO) co-cultured with human T cells and the SLC45A2/PMEL TriTE, we observed a greater cytotoxic response as compared to DMSO-treated cells incubated with the PMEL or SLC45A2 BiTEs alone (**Fig. 6H, Fig. S11E**), reducing the IC_50_ to 0.7 nM (**Fig. 6L**). Overall, we demonstrate T-cell engagers against TAA pMHC’s can induce cytotoxicity in melanoma and in several cases, MEKi-treated melanoma lines can enhance this cytotoxic effect, thus potentially providing a therapeutic strategy using a combination of MEKi and targeted immunotherapies. Future work may focus on further characterizing the pMHC-Abs highlighted here and evaluating the potential for enhanced cytotoxicity in combination with MEKi treatment in an *in vivo* system across clinically relevant treatment timelines and dosing concentrations.

## DISCUSSION

The emergence of drug resistance and/or toxicities to small molecule targeted therapies and checkpoint immunotherapies remains a significant barrier to achieving complete remission in BRAF or NRAS mutant melanoma. To better understand how to optimally combine MEKi with immunotherapy, here we performed a comprehensive analysis characterizing pMHC repertoire response to MEK inhibition using relative and absolute quantitative immunopeptidomics. We identify significantly enriched TAA presentation as a common mechanism to MAPK pathway inhibition *in vitro* & *in vivo* in NRAS-mutant and BRAF-mutant melanomas. While elevated surface HLA presentation in response to MEKi has been previously reported, our data reveal that many of the enriched TAAs increased well beyond average changes in HLA surface expression, in some cases more than 10-fold.

Elevated TAA presentation was observed across varying levels of sensitivities to MEKi, including at sub-cytotoxic doses, and was common to NRAS-mutant and BRAF-mutant lines, suggesting that this response may be shared across many melanoma patients. A multi-omics analysis highlighted changes in cellular plasticity following MEKi as a likely mechanism for the upregulation of certain TAAs. This finding was supported by patient data from the TCGA and suggests that MAPK pathway inhibition may selectively enhance presentation of shared tumor antigens, particularly melanoma differentiation antigens, making them attractive therapeutic targets.

One of the primary criticisms of utilizing shared tumor associated/tissue differentiation antigens (as opposed to tumor-specific antigens, i.e. “neoantigens”) as a therapeutic target for TCR-based therapies is that their low expression in non-tumor tissue can lead to off-target toxicity, likely attributed to the high sensitivity of T cells (45-47). We hypothesized these antigens with high basal or MEKi-induced expression could be intelligently leveraged using antibody-based therapies, which require higher thresholds of antigen presentation for efficacy than TCR-focused approaches, limiting off target toxicity in low-expressing, non-target tissue.

Here, four pMHC-specific antibodies were generated and incorporated into ADC and BiTE formats. Using these reagents, we demonstrated enhanced cell killing following MEKi treatment with either therapeutic modality. Cytotoxicity is observed using just the endogenous or MEKi-augmented antigen presentation levels, in contrast to engineered or overexpression cellular system, which may be less likely to represent physiologically relevant epitope densities (26). Importantly, this work connects targeted immunotherapy response to epitope abundance measurements made using embedded hipMHC multipoint calibrants for accurate quantitative estimations. This is distinct from studies employing exogenous peptide standards for absolute quantification, which underestimate copy-per-cell estimations due to significant losses occurring during sample processing, leading to inaccurate conclusions regarding the sensitivity profile of a given pMHC-targeted modality (25, 48).

Here, we confirm that high (>4e4 copies-per-cell) surface expression is required for ADC efficacy, though this high-level expression is rare and therefore not an optimal strategy for a majority of pMHC epitopes. In contrast, BiTES were effective at lower epitope densities, where the greatest difference in cytotoxic response was observed when cells had fewer than ~1000 copies-per-cell. BiTES showed similar efficacy against targets present at 1000 copies-per-cell or higher, though future studies exploring more pMHCs may further elucidate the relationship between antibody affinity and epitope density. While MEKi treatment did not augment HLA presentation levels in primary melanocytes, future studies may apply SureQuant-IsoMHC to estimate TAA epitope abundances in non-malignant cell lines or primary tissues to better understand the potential for off-target toxicity (49).

In this study we primarily tested the cytotoxicity of a single pMHC-specific BiTE on tumor cells, yet BiTEs could be used in combination to enhance efficacy, or engineered as TriTEs against different epitopes for a single TAA or two different TAAs, offering a multitude of “off the shelf” targeted immunotherapy opportunities to target highly abundant, shared TAAs. Peptide MHC-specific antibodies and MEKi-induced expression could also be utilized for other antibody-based therapeutic strategies such as to initiate antibody-mediated cellular cytotoxicity (ADCC) (41), fabs conjugated to immunotoxins (33), or engineered as pMHC-specific chimeric antigen receptor T-cells (50), where higher expression may also enhance efficacy and/or improve the therapeutic window. Furthermore, although the focus of the therapeutic modalities generated in this study was limited to HLA-A2:01, the same strategy could be employed for other high frequency alleles using MEKi-modulated TAAs identified within this study.

Though resistance to MEKi is inevitable for many melanoma patients, utilizing MEKi to boost TAA antigen presentation prior to or concurrently with ICI and antigen-specific immunotherapies like those described within this study or others (ex. vaccines, cell therapy) may improve therapeutic response. Beyond melanoma, a variety of different therapeutic modalities across cancer types have also been demonstrated to enhance HLA presentation (51-53). Employing quantitative immunopeptidomics in these settings may unlock additional treatment-modulated tumor antigens and provide critical insights as to how to appropriately leverage them for optimal therapeutic potential.

## METHODS

### Human cell lines

SKMEL5, SKMEL28, A375, RPMI-7951, and T2 cell lines were obtained from ATCC [ATCC HTB-70, ATCC HTB-72, CRL1619, HTB-66, and CRL-1992 respectively] and maintained in DMEM medium (Corning). IPC298 and SKMEL2 cells were provided by Array Biopharma and maintained in RPMI 1640 (Gibco) and MEM-α (Gibco) mediums, respectively. Primary epidermal melanocytes (normal, human, adult) were obtained from ATCC (PCS-200-013) and maintained in dermal cell basal medium (ATCC PCS-200-030) supplemented with adult melanocyte growth kit (ATCC PCD-200-042). NFAT-GFP Jurkat cells were a generous gift from Dr. Arthur Weiss (UCSF) and were maintained in RPMI1640 + 2 mg/mL Geneticin (Gibco). All medium was supplemented with 10% FBS (Gibco) and 1% penicillin/streptomycin (p/s, Gibco) except for primary melanocytes (p/s only). Cells were routinely tested for mycoplasma contamination, and maintained in 37 °C, 5% CO_2_. All experiments were performed on passages 4-10.

### Cell line xenografts

SKMEL5, SKMEL28, SKMEL2, and IPC298 cell lines were used for cell line xenograft (CLX) analyses in collaboration with Array Biopharma. 5×10^6^ cells in 100 μL phosphoate buffered saline (PBS) containing 50% Matrigel^®^ were implanted via subcutaneous injection into NCr nu/nu mice on the right flank. Resultant tumors were randomized into study groups at a starting size of ~200-400mg, dosed at 10 mL/kg for up to five days by oral gavage with vehicle, 3.5 mg/kg binimetinib (MEK162) or 20 mg/kg encorafenib (LGX818) prepared as suspensions in 1% CMC/ 0.5% Tween 80. Dosing continued for up to 5 days, and at the end of each time course tumors were harvested and flash frozen in liquid nitrogen. Animals were housed in groups of 3. Food, water, temperature, and humidity are according to Pharmacology Testing Facility performance standards (SOP’s) which are in accordance with the 2011 Guide for the Care and Use of Laboratory Animals (NRC) and AAALAC-International. Dosing schedules are listed in **SI Appendix, Table S4**.

### Peptide synthesis

Heavy leucine-containing peptides for hipMHC quantification correction (ALNEQIARL^+7^, SLPEEIGHL^+7^, and SVVESVKFL^+7^ were synthesized at the MIT-Koch Institute Swanson Biotechnology Center in Biopolymers and Proteomics Facility using standard Fmoc chemistry using an Intavis model MultiPep peptide synthesizer with HATU activation and 5 μmol chemistry cycles as previously described.(24) Standard Fmoc amino acids were procured from NovaBiochem and Fmoc-Leu (13C6, 15N) was obtained from Cambridge Isotope Laboratories. Light peptides for pMHC-antibody generation (PMEL, DCT, PRUNE2, SLC45A2) were synthesized on a Gyros-Protein Technologies Tribute with UV feedback at a 100 micromole scale using standard Fmoc chemistry and HATU/NMM activation chemistry. Both light and heavy leucine-containing peptides were purified on a Gilson GX-271 preparative HPLC system by reverse phase, and quality assured with MS on a Bruker MicroFlex MALDI-TOF and by RP-HPLC on an Agilent model 1100 HPLC.

Isotopologue peptides for SureQuant-IsoMHC analyses were synthesized using HeavyPeptide AQUA Custom Synthesis Service (Thermo Fisher Scientific) and were purified to >97% and validated with amino acid analysis as previously described (40).

### Cloning

Fabs were subcloned from the Fab-phagemid into an *E. coli* expression vector pBL347. The heavy chain of the IgG was cloned from the Fab plasmid into a pFUSE (InvivoGen) vector with a human IgG1 Fc domain. The light chain of the IgG was cloned from the Fab plasmid into the same vector but lacking the Fc domain. The light chain of the BiTE was cloned from the Fab plasmid into a pFUSE (InvivoGen) vector with an anti-CD3 scFv (OKT3). The heavy chain of the BiTE was cloned into the same vector lacking the OKT3. SCD3-arm of the TriTE was converted into a scFab and cloned into a pFUSE (InvivoGen) vector with the KIH strategy “knob” human Fc domain(54) MLA2-arm of the TriTE was converted into a scFab and cloned into a pFUSE (InvivoGen) vector with the KIH strategy “hole” human Fc domain followed by OKT3. All constructs were sequence verified by Sanger sequencing.

### Protein expression and purification

MHC-peptide complexes were expressed and refolded as previously described.(55) Briefly, MHC-peptide complexes were refolded at 10°C for 3 days and SEC-purified on a HiLoad 16/600 Superdex 75 pg column equilibrated in 10 mM Tris pH 8. After purification, MHC-peptide complexes were biotinylated using a BirA reaction kit (Avidity) per manufacturer’s instructions in the presence of excess peptide and β_2_M at 25°C for 4 hours. After biotinylation, MHC-peptide complexes were purified again via SEC to remove excess biotin. Proper folding was assessed by SDS-PAGE. Biotinylation was assessed by preincubating MHC-peptide complexes with NeutrAvidin and subsequently assessed by SDS-PAGE.

Fabs were expressed in E. coli C43 (DE3) Pro+ as previously described using an optimized autoinduction medium and purified by protein A affinity chromatography (56). IgGs, BiTEs, and TriTEs were expressed in Expi293 BirA cells using transient transfection (Expifectamine, Thermo Fisher Scientific). After transfection for 3-5 d, media was harvested, IgGs and TriTEs purified by Ni-NTA affinity chromatography and BiTEs were purified using protein A affinity chromatography. All proteins were buffer exchanged into PBS pH 7.4 and stored in 10% glycerol at −80°C and assessed by SDS-PAGE.

All proteins were then buffer exchanged into phosphate-buffered saline (PBS) containing 20% glycerol, concentrated, and flash frozen for storage. All other proteins were buffer exchanged into PBS by spin concentration and stored in aliquots at −80°C. The purity and integrity of all proteins were assessed by SDS-PAGE. Fabs were subsequently buffer exchanged into PBS pH 7.4 and stored in 10% glycerol at −80°C and assessed by SDS-PAGE.

### Fab-phage selection

Phage selections were run as previously described (Hornsby et al. 2015). Selections were performed on a KingFischer™ System (Thermo Fisher Scientific). Biotinylated antigens were immobilized using streptavidin-coated magnetic beads (Promega). In each round, phage was first cleared by incubation with beads loaded with MHC-peptide complexes loaded with FLU peptide. Unbound phage was next incubated with beads loaded with MHC-peptide complex of interest. Beads were washed and bound phage was eluted with 50 μg/mL of TEV protease. Four rounds of selection were performed with decreasing amounts of MHC-peptide complex of interest. Selections were performed in PBS+0.02% Tween-20+0.2% bovine serum albumin (PBSTB). Individual phage clones from the fourth round of selections were analyzed by ELISA.

### Phage ELISA

For each phage clone, four different conditions were tested - Direct: MHC-peptide complex of interest, Competition: MHC-peptide complex of interest with an equal concentration of MHC-peptide complex in solution, Negative selection: FLU MHC-peptide complex, and Control: PBSTB. 384-well Nunc Maxisorp flat-bottom clear plates (Thermo Fisher Scientific) were coated with 0.5 μg/mL of NeutrAvidin in PBS overnight at 4°C and subsequently blocked with PBSTB. Plates were washed 3x with PBS containing 0.05% Tween-20 (PBST) and were washed similarly between each of the steps. 20 nM biotinylated MHC-peptide complex was diluted in PBSTB and immobilized on the NeutrAvidin-coated wells for 30 minutes at room temperature, then blocked with PBSTB + 10 μM biotin for 10 minutes. For the competition samples, phage supernatant was diluted 1:5 into PBSTB with 20 nM MHC-peptide complex of interest for 30 minutes prior to addition to the plate. For the direct samples, phage supernatant was diluted 1:5 in PBSTB. Competition and direct samples were added to the plate for 30 minutes at room temperature. Bound phage was detected by incubation with anti-M13-horseradish peroxidase conjugate (Sino Biologics, 1:5000) for 30 minutes, followed by the addition of TMB substrate (VWR International). The reaction was quenched with the addition of 1 M phosphoric acid and the absorbance at 450 nm was measured using a Tecan M200 Pro spectrophotometer. Clones with high binding to MHC-peptide complex of interest, low binding to PBSTB/FLU MHC-peptide complex, and a competition ratio (Competition AU/Direct AU) ≥0.5 were carried forward.

### Bio-layer Interferometry

BLI measurements were made using an Octet RED384 (ForteBio) instrument. MHC-peptide complex was immobilized on an streptavidin biosensor and loaded for 200 seconds. After blocking with 10 μM biotin, purified binders in solution were used as the analyte. PBSTB was used for all buffers. Data were analyzed using the ForteBio Octet analysis software and kinetic parameters were determined using a 1:1 monovalent binding model.

### IgG NHS-Fluorophore Conjugation

Purified IgG’s were buffer exchanged into PBS pH 8.3. Concentrated IgG to ~11 mg/mL (with the exception of P2B1 which was only 2 mg/mL), and added 20 mM NHS-AF488 (Fluoroprobes) at either a 10:1 or 5:1 (Dye:IgG) ratio. Conjugation reactions were incubated at room temperature for 1 hour, and then quench by adding equivalent volume of 1 M glycine pH 8.4 as dye. Reactions were further incubated for 1 hour and then buffer exchanged into PBS pH 7.4 until all excess dye was removed. IgG and dye concentration was determined by UV.

### ADC conjugation

Purified IgG was buffer exchanged into PBS pH 7.4 and concentrated to 35μM. 20x 100 mM piperidine-derived oxaziridine molecule (57) was added to PBS pH 7.4, and subsequently added to IgG for a final IgG concentration of 35μM. Labeling was conducted at room temperature for 2 hours, and buffer exchanged with PBS pH 7.4 to remove unconjugated oxaziridine. 5% v/v 5 mM DBCO-PEG4-Glu-vc-PAB-MMAF (Levena Biopharma) was added to oxaziridine-labeled IgG and incubated overnight at room temperature. IgG was buffer exchanged into PBS pH 7.4 to remove unconjugated MMAF. Conjugation efficiency was assessed by intact protein mass spectrometry using a Xevo G2-XS Mass Spectrometer (Waters).

### Flow cytometry

#### Surface HLA expression in melanoma cells

Cells were seeded and treated with DMSO or binimetinib in 10 cm plates, then lifted with 0.05% Trypsin-EDTA and 10^6^ cells/mL were spun at 300 g for 3 minutes, washed with ice cold flow buffer [1X PBS supplemented with 3% bovine serum albumin (BSA)] and incubated with fluorophore-conjugated antibody at 0.5 μg mL^-1^ in flow buffer for 30 minutes on ice. After incubation, cells were washed again, and resuspended in flow buffer plus 5 μL of propidium iodide (PI) staining solution (10 μg mL^-1^, Invitrogen) per sample. Analyses were performed on an LSRII (BD Biosciences) and all flow cytometry data was analyzed using FlowJo (version 10.7.2). Antibody: Alexa Fluor 488 HLA-A, B, C, clone W6/32 [Biolegend, cat # 311413]. The gating strategy previously described (24).

#### pMHC-Fab and pMHC-antibody staining

T2 lymphoblasts: the day prior to Fab staining, T2 lymphoblasts were cultured in RPMI serum-free media containing 50 μg/mL peptide of interest at a concentration of 1e^6^ cells/mL. Cells were collected by centrifugation and washed 1X in flow buffer. Each sample was resuspended in 10 μg/mL Fab for 30 minutes, and then washed 3x in flow buffer. Each sample was then stained with an anti-human Fab goat mAb Alexa Fluor 647 conjugate (Jackson ImmunoResearch) for 30 minutes, and then washed 3x in flow buffer. Samples were resuspended in 200 μL sterile PBS pH 7.4 and analyzed on a CytoFLEX (Beckman Coulter). For pMHC-antibody staining (full length IgG), T2 cells were incubated with the peptide of interest overnight, harvested, and stained with either primary pMHC specific IgG antibodies or a human IgG isotype control (Abcam, ab20619) at 10 *μ*g/mL for 20 minutes on ice. Cells were then washed with flow buffer 1X and incubated with protein A-488 secondary antibody conjugate (Invitrogen, P11047) for 20 minutes (1:1000 dilution). Cells were washed again with flow buffer and resuspended in PI staining solution prior to analysis on the LSRII. Gating strategy for T2 cells is shown in **Fig. S12A**.

SKMEL5 cells: SKMEL5 cells were pre-treated with DMSO or 1 *μ*M binimetinib for 72 hours in 10 cm plates and were subsequently harvested (10^6^ cells/mL), washed, and stained with fluorophore conjugated pMHC-antibodies at 2 *μ*g/mL, and analyzed using the LSRII. **Fig. S12B** describes the gating strategy for SKMEL5 cells.

#### Jurkat NFAT-GFP activation

SKMEL5 cells were treated in 10 cm plates with DMSO or 1 *μ*M binimetinib for 72 hours, after which cells were seeded in a 24 well plate at a ratio of 250,000 SKMEL5 cells to 50,000 Jurkat NFAT-GFP cells (5:1) in Jurkat culture medium with n=3 technical replicates per condition and incubated with a pMHC-specific or anti-GFP (control) BiTE for 24 hours. T2 cells were seeded at 1:1 ratio (5e^4^ cells to 5e^4^ cells) in a 96-well round bottom plate. Cells were washed 2x with flow buffer and resuspended in PI staining solution. Cells were gated according to the stagey described in **Fig. S12C**, where the percentage of GFP positive cells were gated so ~97% of Jurkat cells with no BiTE were classified as GFP negative. SKMEL5 cells were analyzed on the LSRII, T2 on the CytoFLEX.

### Cell viability assays

#### Binimetinib dose response

Half-maximal inhibitory concentrations (IC_50_) of binimetinib (Selleckchem, MEK162) were determined for each cell line using CellTiter-Glo (CTG) luminescent cell viability assay (Promega). Cells were seeded at density of 10,000 (SKMEL2, SKMEL28, IPC298) or 5,000 (SKMEL5, A375, RPMI-7951) cells/well in a 96 well plate and allowed to adhere overnight. Cells were then treated with binimetinib or DMSO as a vehicle control in fresh medium for 72h and assayed. All viability data was acquired using a Tecan plate reader Infinite 200 with Tecan icontrol version 1.7.1.12. IC_50_ values were calculated using a 4-parameter logistic curve in Prism 9.0.0.

#### Antibody-drug conjugate cell killing assays

SKMEL5 or A375 cells were pre-treated for 72 hours with DMSO or 1 *μ*M binimetinib in 10 cm plates, and subsequently seeded at a density of 5,000 cells/well in a 96 plate. Cells were incubated antibody-drug conjugate with n=4 technical replicates per treatment condition for an additional 72 hours and similarly assayed with CTG.

#### T-cell/target cell co-incubation cell killing assays

Deidentified buffy coats from healthy human donors were obtained from Massachusetts General Hospital. Peripheral blood mononuclear cells (PBMCs) were isolated by density-based centrifugation using Ficoll (GE Healthcare). CD8+ T cells were isolated from PBMCs using a CD8+ T cell negative selection kit (Stemcell). T cells were mixed with Human T-activator CD3/CD28 DynaBeads (Thermo Fisher Scientific) in a 1:1 ratio and maintained in R10 + IL-2 [RPMI 1640 (Thermo Fisher Scientific) supplemented with 10% heat-inactivated FBS (Thermo Fisher Scientific), 1% HEPES (Corning), 1% L-glutamine (Thermo Fisher Scientific), 1% Pen/Strep (Corning) and 50 IU/mL of IL-2 (R&D Systems)] for 7 days prior to use in cell killing assays. DynaBeads were removed by magnetic separation prior to coincubation of primary T cells with target cells. Target cells were treated with DMSO or 1 *μ*M MEKi for 72 hours and were subsequently seeded in a 96-well plate with primary T-cells in R10 + IL-2 at an effector to target ratio of 2:1 and incubated with BiTEs for 48 hours with n=3 technical replicates per condition. Cells were assayed with CTG, and percent cytotoxicity was calculated by subtracting the average luminescence signal of the T-cell only condition and normalizing to the no BiTE condition. ((X-[T-cell only]) / ([average-no-BiTE] -[T-cell only])) x 100.

### UV-mediated peptide exchange for hipMHCs

UV-mediated peptide exchange was performed using recombinant, biotinylated Flex-T HLA-A*02:01 monomers (BioLegend), using a modified version of the commercial protocol. Briefly, 2-4 μL of 500 μM peptide stock, 2 μL of Flex-T monomer, and 32 μL of 1X PBS were combined in a 96-well U bottom plate. On ice, plates were illuminated with ultraviolet light (365 nm) for 30 minutes, followed by a 30-minute incubation at 37 °C protected from light. Concentration of stable complexes following peptide exchange was quantified using the Flex-T HLA class I ELISA assay (Biolegend) per manufacturer’s instructions for HLA-A*02:01. ELISA results were acquired using a Tecan plate reader Infinite 200 with Tecan icontrol version 1.7.1.12.

### Peptide MHC isolation

Cultured cells were seeded in 10 cm plates, allowed to adhere overnight, and treated for 72h with binimetinib or DMSO vehicle control. At the time of harvest, cells were washed with 1X PBS, and lifted using 0.05% Trypsin-EDTA (Gibco). Cells were pelleted at 500 g for 5 minutes, washed twice more in 1X PBS, and pelleted again. Cells were resuspended in 1 mL lysis buffer [20 nM Tris-HCl pH 8.0, 150 mM NaCl, 0.2 mM PMSO, 1% CHAPS, and 1X HALT Protease/Phosphatase Inhibitor Cocktail (Thermo Scientific)], followed by brief sonication (3 x 10 second microtip sonicator pulses) to disrupt cell membranes. Lysate was cleared by centrifugation at 5000 g for 5 minutes and quantified using bicinchoninic acid protein assay kit (Pierce). For *in vitro* analyses, 1×10^7^ cells were used for each condition. Frozen CLX tumor samples were homogenized in lysis buffer, cleared by centrifugation, and quantified using BCA as described in the in vitro analyses. For each sample, 7 mg of lysate was used. For absolute quantification analyses, ~5 mg of lysate was used.

Peptide MHCs were isolated by immunoprecipitation (IP) and size exclusion filtration, as previously described.(24) Briefly, 0.5 mg of pan-specific anti-human MHC Class I (HLA-A, HLA-B, HLA-C) antibody (clone W6/32, Bio X Cell [cat # BE0079]) was bound to 20 μL FastFlow Protein A Sepharose bead slurry (GE Healthcare) for 3 hours rotating at 4 °C. Beads were washed 2x with IP buffer (20 nM Tris-HCl pH 8.0, 150 mM NaCl) prior to lysate and hipMHC addition (*in vitro analyses*), and incubated rotating overnight at 4 °C to isolate pMHCs. For TMT-labeled DDA analyses, 30 fmol of the following hipMHC standards were added prior to IP for quantification correction: ALNEQIARL^7^, SLPEEIGHL^7^, and SVVESVKFL^7^. For absolute quantification analyses, 1, 10, or 100 fmol of 1-3H Iso18 hipMHCs standards were added to each immunoprecipitation. Beads were washed with 1X Tris buffered saline (TBS) and water, and pMHCs were eluted in 10% formic acid for 20 minutes at room temperature (RT). Peptides were isolated from antibody and MHC molecules using a passivated 10K molecule weight cutoff filter (PALL Life Science), lyophilized, and stored at −80 °C. Label-free MS analysis acquisition parameters and data analysis techniques are described in the **SI Appendix, Methods**.

### pMHC labeling with Tandem Mass Tags and SP3 cleanup

For labeled analyses, 100 μg of pre-aliquoted Tandem Mass Tag (TMT) 6-plex, 10-plex, or TMT-pro was resuspended in 30 μL anhydrous acetonitrile, and lyophilized peptides were resuspended in 100 μL 150 mM triethylammonium bicarbonate, 50% ethanol. Both were gently vortexed, centrifuged at 13,400 g for 1 minute, and combined. TMT/peptide mixtures were incubated on a shaker for 1 hour at RT, followed by 15 minutes of vacuum centrifugation. After combining labeled samples, we washed tubes 2x with 25% acetonitrile (MeCN) in 0.1% acetic acid (AcOH) and added it to the labeled mixture, which was subsequently centrifuged to dryness.

Sample cleanup was performed using single-pot solid-phase-enhanced sample preparation (SP3) as previously described.(58) Briefly, a 1:1 mix of hydrophobic/hydrophilic Sera-mag carboxylate-modified speed beads (GE Healthcare) was prepared at a final bead concentration of 10 μg μL^-1^. Labeled samples were resuspended in 30 μL of 100 mM ammonium bicarbonate (pH 7-8) and added to 500 μg of bead mix with 1 mL MeCN. Peptides were allowed to bind for 10 minutes at RT, washed 2x with MeCN, and eluted with 2% DMSO for 1 minute of sonication in a bath sonicator. TMT-labeled peptides were transferred to a fresh microcentrifuge tube and centrifuged to dryness. Peptides were resuspended in 0.1% formic acid, 5% MeCN and analyzed by MS. MS acquisition parameters and data analysis techniques are described in the **SI Appendix, Methods**.

### Global protein expression profiling sample preparation

For a quantitative global proteomics analysis, 300 μg of supernatant from DMSO and 100 nM MEKi sample for SKMEL5 cells was diluted 8-fold in 8M urea, reduced with 10 mM dithiothreitol in 100 mM ammonium acetate (pH 8.9) at 56°C for 45 minutes, and subsequently alkylated with 50 mM iodoacetamide for 45 minutes rotating at RT in the dark. Lysates were diluted 4-fold with 100 mM ammonium acetate and digested with sequence-grade trypsin (Promega) overnight at RT at an enzyme:substrate ratio of 50:1 (w/w). The reaction was quenched with formic acid (5% total volume) and desalted on C18-based STAGE tips. Solvents: 0.1% formic acid, 90% acetonitrile (MeCN) in 0.1% formic acid, and 60% acetic MeCN in 0.1% formic acid. Volumes were reduced with vacuum centrifugation and lyophilized in 150 ug aliquots. Peptide aliquots were labeled with TMT10-plex reagents in 70% ethanol/150 mM triethylammonium bicarbonate (TEAB) for 1 hour at room temperature, pooled, brought to dryness with vacuum centrifugation, and stored at −80°C.

The labeled mixture was resuspended in 0.1% formic acid, and 25% was loaded onto an Agilent Zorbax 300Extend-C18 5 μm 4.6 × 250 mm column on an Agilent 1200 operating at 1 ml/min for fractionation, as previously described (59). Briefly, peptides were eluted with the following gradient: 1% B to 5% B for 10 mins, 5-35% B for 60 mins, 35-70% B for 15 min, held at 70% B for 5 mins, and was followed by equilibration back to 1% B. Fractions were collected with a Gilson FC203B fraction collector at 1 minute intervals and fractions 10-90 were concatenated to 20 fractions. The fraction volumes were next reduced by vacuum centrifugation, lyophilized, and stored at −80°C prior to analysis. MS acquisition parameters and data analysis techniques are described in the **SI Appendix, Methods**.

### Ubiquitination sample preparation

SKMEL5 cells were seeded in 10 cm plates and allowed to adhere overnight. Cells were then treated with DMSO or 100 nM binimetinib for 72 hours. Prior to harvest, cells were treated with 100 nM bortezomib (PS-341, SelleckChem) to halt protease activity. Cells were next washed with ice cold 1X PBS and lysed in 8M Urea. Lysates were processed to tryptic peptides as described in the global protein expression methods and desalted using SepPak plus cartridges. Five mg aliquots per sample were lyophilized and stored at 80 °C prior to analysis.

PRMScan ubiquitin remnant motif (anti-K-ε-GG) antibody beads (Cell Signaling Technology, #5562) were crosslinked as previously described.(60) Briefly, beads were washed 3x with 100 mM sodium borate pH 9, incubated in cross linking buffer (20 mM DMP in 100 mM sodium borate pH9) for 30 mins (RT, rotation). Beads were next washed 3x with blocking buffer (200 mM ethanolamine, pH 8) and incubated for 2 hours at 4°C rotating. Crosslinked beads were washed 3x with immunoprecipitation buffer (100 mmol/l Tris-HCl, 1% Nonidet P-40 at pH 7.4) and stored in 1X PBS with 0.02% sodium azide at 4°C prior to use.

Each sample was resuspended in 1 mL IP buffer and added to 40 uL bead slurry of conjugated anti-K-ε-GG beads and incubated for 2 hours rotating at 4°C.(61) Peptides were washed 2x with IP buffer and 3x with 1X PBS, and diGly peptides were eluted 2x with 0.2% TFA for 5 minutes. To improve specificity, each lysate was IP’d twice, following the same IP protocol with the first elution. Finally, peptides were dried with vacuum centrifugation and lyophilized.

Lyophilized samples were next labeled with 100 μg of TMT-6plex, as described in the MHC labeling methods section. A high pH reverse-phase peptide fraction kit was used to separate labeled peptides into six fractions, according to manufacturer’s instructions (17.5%, 20%, 22.5%, 25%, 30%, and 70% MeCN, Thermo Scientific). Peptide fraction volume was reduced with vacuum centrifugation, lyophilized, and stored at −80°C prior to analysis. MS acquisition parameters and data analysis techniques are described in the **SI Appendix, Methods**.

### RNA-sequencing

RNA was isolated from 10 cm plates of SKMEL5 cells with 3 biological replicates per condition (DMSO, 100 nM binimetinib, 1 *μ*M binimetinib) using Direct-zol RNA miniprep kit (Zymo Research), as previously described (24). RNA were confirmed for quality using the Agilent Fragment Analyzer and 300 ng of material was polyA-selected using NEBNext Poly(A) mRNA Magnetic Isolation Module (E7490) modified to include two rounds of polyA binding and 10 minute incubations. cDNA was generated using the NEB Ultra II directional kit (E7760) following manufacturer instructions using 12 cycles of PCR and and a 0.9X SPRI clean. The resulting libraries were quality assessed using the Fragment Analyzer and quantified by qPCR prior to be sequenced on the Illumina HiSeq2000. The 40nt single-end reads with an average depth of 5 million reads per sample were sequenced for all conditions.

RNAseq reads were aligned to the human transcriptome prepared with the hg38 primary assembly and the Ensembl version 95 annotation using STAR version 2.5.3a (62). Gene expression was summarized with RSEM version 1.3.0 and SAMtools version 1.3.(63, 64) Differential expression analysis was performed with DESeq2 version 1.24.0 running under R version 3.6.0 with normal log fold change shrinkage (65). Significance values (adjusted p-value, Wald test) were multiple hypothesis corrected using Benjamini-Hochberg (BH) method. The resulting data were parsed and assembled using Tibco Spotfire Analyst version 7.11.1.

### Peptide MHC binding affinity

Binding affinity of 9-mer pMHCs was estimated using NetMHCpan-4.0 against each cell line’s allelic profile (SI Appendix, **Table S1**) (66, 67). The minimum predicted affinity (nM) of each peptide was used to assign peptides to their best predicted allele. The threshold for binding was set to 500 nM.

### Enrichment analyses

For pMHC pathway and TAA enrichment analyses, gene names from peptide source proteins were extracted and rank ordered according to the average log2 fold change over DMSO treated cells. In cases where more than one peptide mapped to the same source protein, the maximum/minimum was chosen, depending on the directionality of enrichment analysis. For RNAseq & protein expression data, data sets were rank ordered according to the mean log2 fold change value with only protein encoding genes considered.

We utilized gene set enrichment analysis (GSEA) 4.0.3 pre-ranked tool against the Molecular Signatures Database hallmarks gene sets with 1000 permutations, weighted enrichment statistic (p=1), and a minimum gene size of 15 (35, 68, 69). Results were filtered for FDR q-value ≤ 0.25, and nominal p-value ≤ 0.05. P-values > 0.05 in reported analyses are noted.

### TCGA/gTEX/Cell line expression analysis

mRNASeq normalized gene expression data (MD5) from the TCGA skin cutaneous melanoma study (SKCM) for was obtained from Firebrowse (39). Expression for all tumors was z-score normalized, and *BRAF* mutant tumor data was extracted for subsequent analyses. Pairwise gene expression significance comparisons were calculated using an un-paired, two-tailed T test, and significance values for HLA expression between MITF-low and immune subtypes were calculated using Sidak’s multiple comparisons test. Tumor versus normal expression profiles for SKCM were generated using the Gene Expression Profiling Interactive Analysis 2 (GEPIA2) (70) using data from TCGA and gTEX studies with a jitter size of 0.4 and a p-value cutoff of 0.01 for significance, calculated using a one-way ANOVA statistical test. Expression data for 57 BRAF/NRAS melanoma cell lines was obtained from TRON Cell line Portal (67).

## Data availability

RNA-sequencing data have been deposited into the NCBI Gene Expression Omnibus GSE85284. The mass spectrometry proteomics data have been deposited to the ProteomeXchange Consortium via the PRIDE partner repository with the dataset identifier PXD029860 for DDA datasets and PXD029884 for targeted datasets.

## ACKNOWLEDGEMENTS

We thank Ryan Sullivan & Genevieve Boland for project guidance and feedback, Alex Jaegar and Connor Dobson for guidance on T-cell cytotoxicity assays, Iris Abrahantes Morales for assistance with T-cell isolation, Susanna Elledge for assistance with bioconjugation of IgG’s, and Aaron Gajadhar, Bhavin Patel, and Sebastien Gallien from Thermo Fisher for project support on SureQuant-MHC. We also thank the MIT BioMicro Center (Stuart Levine) and the Swanson Biotechnology Center for technical support, specifically the Flow Cytometry (Glenn Paradis), Biopolymers & Proteomics (Richard Cook), and the Barbara K. Ostrom (1978) Bioinformatics (Charlie Whittaker) core facilities. This research was supported in part by NIH R35GM122451 and NCI R01CA248323 (J.A.W.), NIH U54 CA210180 and U01CA238720 (F.M.W.), as well as funding from the Melanoma Research Alliance (MRA Team Science Award 565436) and the MIT Center for Precision Cancer Medicine. J.A.W. was supported by generous funding from the Chan Zuckerberg Biohub Investigator Program, the Harry and Dianna Hind Professorship; N.J.R. was supported by the National Science Foundation Graduate Research Fellowship; L.E.S. was supported by an NIH training grant in Environmental Toxicology (T32-ES007020). The authors declare that they have no competing interests.

## SI Appendix

Supplementary Figures & legends

Supplementary Methods

Supplementary Data legends

Supplementary Tables & legends

## SUPPLEMENTARY FIGURE LEGENDS

**Fig. S1** Phenotypic characterization of cellular response to binimetinib.

**Fig. S2** MHC peptide characterization from *in vitro* analyses.

**Fig. S3** Volcano plots of *in vitro* pMHC analyses.

**Fig. S4** TAA pMHC enrichment following binimetinib treatment.

**Fig. S5** TAA enrichment following MEKi treatment *in vivo*.

**Fig. S6** Correlation between pMHC, protein, and transcript expression with MEKi treatment.

**Fig. S7** pMHC presentation changes in EMT-derived epitopes.

**Fig. S8** Characterization of Fab-phage clones.

**Fig. S9** Peptide specificity of pMHC-specific Fabs.

**Fig. S10** Tumor versus normal expression profiles of select TAAs.

**Fig. S11** Characterization of pMHC-specific ADCs and BiTEs *in vitro*.

**Fig. S12** Flow cytometry gating strategies.

## SUPPLEMENTARY DATA LEGENDS

**Dataset S1** In vitro quantitative immunopeptidomics datasets for all cell lines and treatment conditions.

**Dataset S2** In vivo quantitative immunopeptidomics for all cell line xenografts and treatment conditions.

**Dataset S3** Transcript expression of SKMEL5 cells +/- 100 nM and 1 uM binimetinib treatment for 72 hours.

**Dataset S4** Protein expression of SKMEL5 cells +/- 100 nM binimetinib treatment for 72 hours.

**Dataset S5** Peptide ubiquitination levels of SKMEL5 cells +/- 100 nM binimetinib treatment for 72 hours.

**Dataset S6** Absolute quantification of 18 tumor associated antigens

## SUPPLEMENTARY TABLE LEGENDS

**Table S1** Allelic profile of melanoma cell lines.

**Table S2** Tumor associated antigen peptide library for enrichment analyses.

**Table S3** Custom library of tumor associated antigen source proteins.

**Table S4** CLX treatment groups and dosing schedule.

