## Supplementary material for "MEK inhibition enhances presentation of targetable MHC-I tumor antigens in mutant melanomas": SI Appendix

Stopfer & Rettko et al.

#### SI Appendix

*This file contains:*
Supplementary Figures & legends
Supplementary Methods
Supplementary Data legends
Supplementary Tables & legends

### SUPPLEMENTARY FIGURES

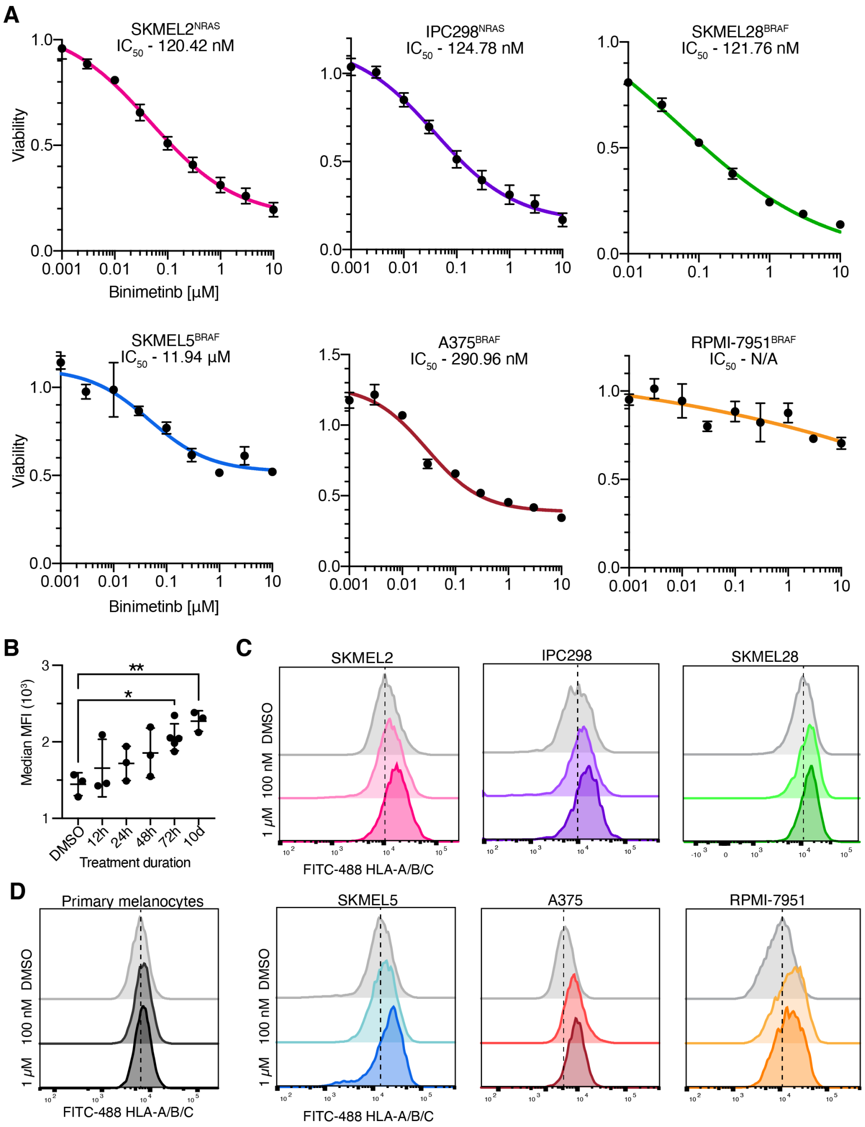


**Fig. S1. Phenotypic characterization of cellular response to binimetinib.**
**(A)** Cell viability (fraction of DMSO control) at 72 hr after binimetinib treatment. Data are represented as mean values +/- SD for n=3 replicates. Lines represents a four-parameter nonlinear regression curve fit. **(B)** Surface HLA expression of SKMEL5 cells treated with 100 nM binimetinib measured by flow cytometry. Data are represented as mean values +/- SD for n=3 replicates and significance between replicates (*p<0.05, **p<0.01) is calculated using a one way ANOVA test comparing each value to the DMSO control. **(C)-(D)** Flow cytometry measurements of surface HLA expression in cells. Data are represented as % of maximum signal, and the distributions are representative of three independent experiments.

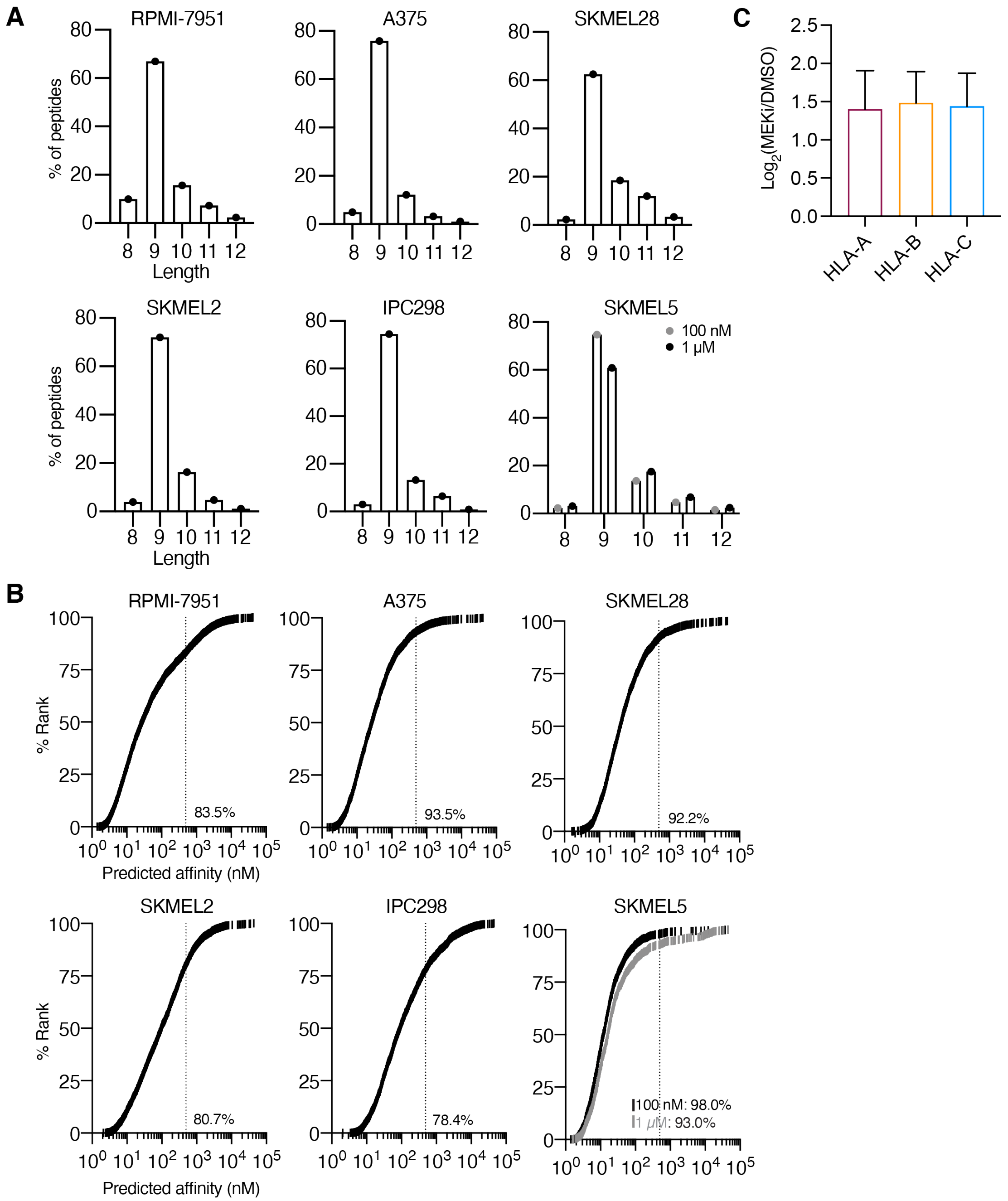


**Fig. S2. MHC peptide characterization from *in vitro* analyses.**

**(A)** Peptide length distribution for each cell line. **(B)** Predicted binding affinity of 9-mer peptides, rank ordered. Dotted line represents threshold for binding at ≤500 nM. Percentage of peptides ≤500 nM are listed on each plot. **(F)** Average fold change in presentation for SKMEL5 9-mers +/- 1 uM MEKi segregated by highest predicted affinity to HLA-A/B/C. Error bars represent +/- stdev. Tukey's multiple comparisons test shows no significant difference between pairwise comparisons.

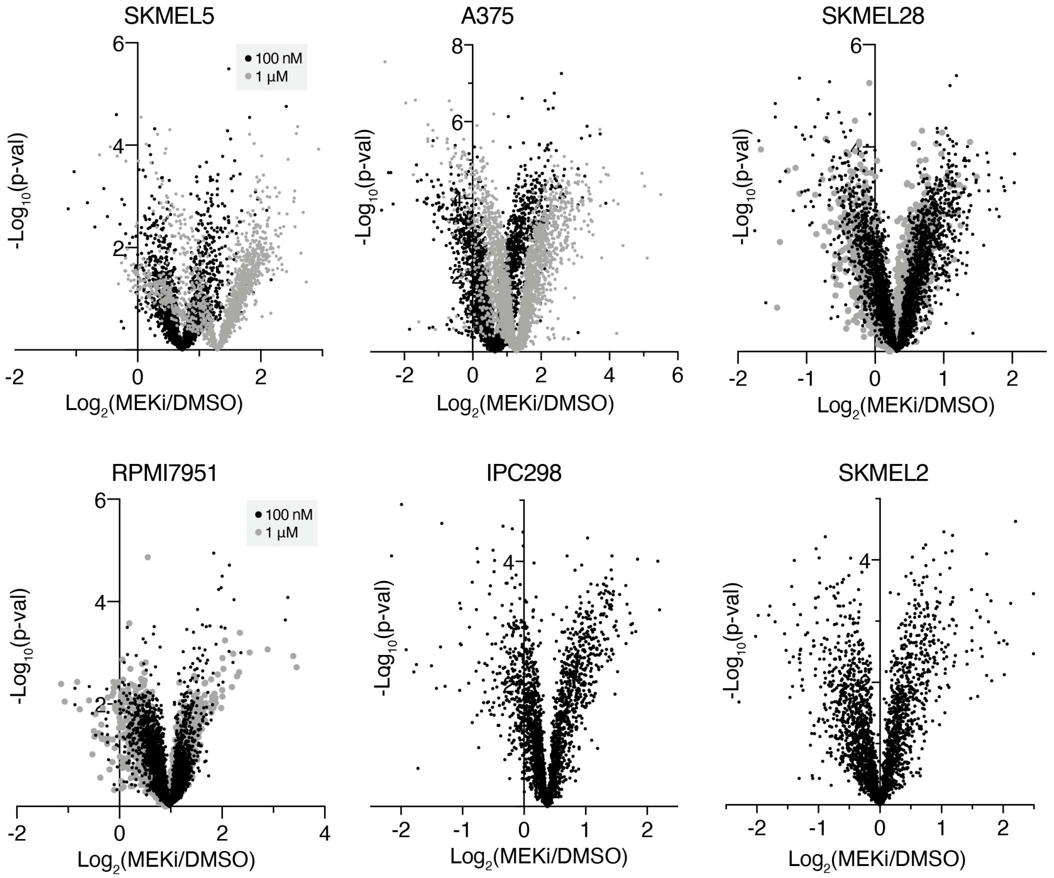


**Fig. S3 Volcano plots of *in vitro* pMHC analyses.**
Volcano plots of the average fold change in pMHC expression with binimetinib treatment (n=3 biological replicates for DMSO and MEKi treated cells) versus significance (mean-adjusted p value, unpaired two-sided t test).

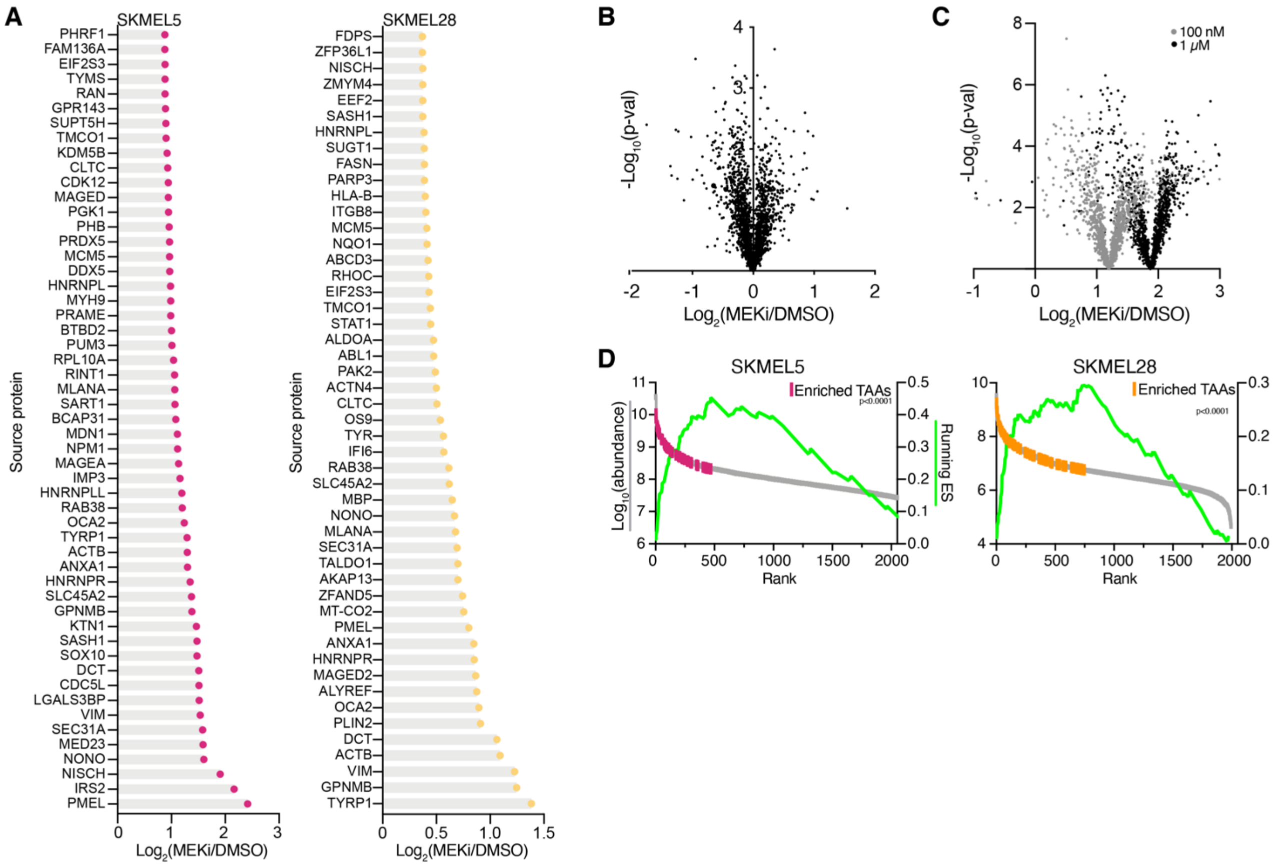


**Fig. S4 TAA pMHC enrichment following binimetinib treatment.**

**(A)** Enriched TAA pMHC expression changes with 100 nM MEKi. **(B)-(C)** Volcano plots of the average fold change in pMHC expression with 10 nM binimetinib treatment **(B)** and 100 nM and 1 µM trametinib treatment **(C)**. Data shown are the mean of n=3 biological replicates per condition versus significance (mean-adjusted p value, unpaired two-sided t test). **(D)** Enrichment plots with peptides rank ordered by precursor ion abundance. p<0.0001 for both SKMEL5 100 nM and SKMEL28 100 nM analyses.

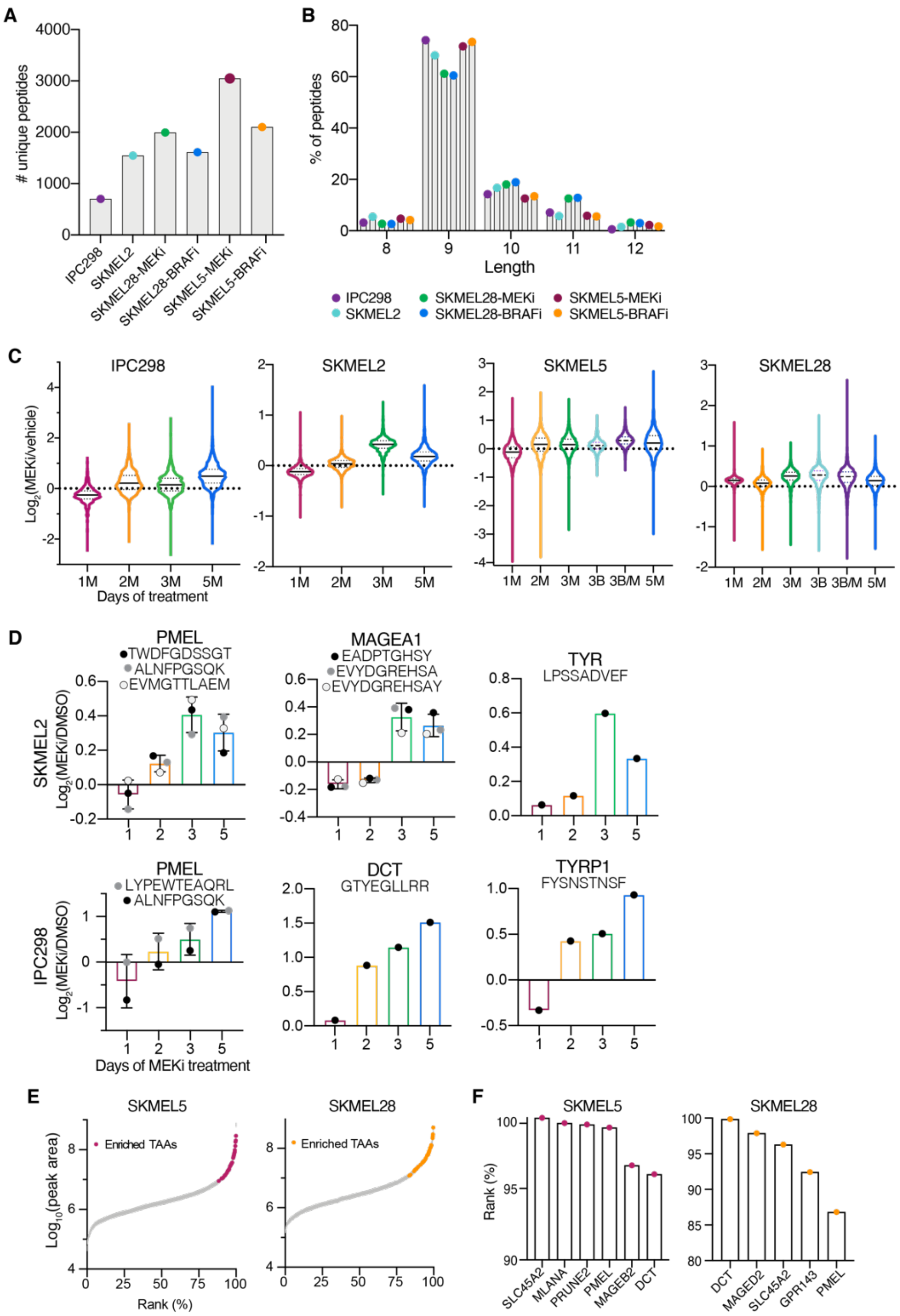


**Fig. S5 TAA enrichment following MEKi treatment *in vivo.***

**(A)** Number of unique pMHCs identified in each analysis. **(B)** Length distribution of pMHCs represented as a percentage of the total pMHCs identified. **C)** Violin plot of distribution of fold changes in presentation of pMHCs following MEK inhibition (M, binimetinib), BRAF inhibition (B, encorafinib) or both (B/M). Solid line represents median, dotted lines define the first and third quartiles. **(D)** Changes in pMHC expression for select melanoma differentiation antigens, x-axis = number of days of binimetinib treatment. Errors bars represent standard deviation when >1 peptide from each source protein was identified. **(E)** Rank-ordered average abundance (n=3 biological replicates) of pMHCs in SKMEL5 and SKMEL28 analyses. Positively enriched TAAs are highlighted in color. **(F)** Select TAA peptide rank-ordered average abundance for CLXs.

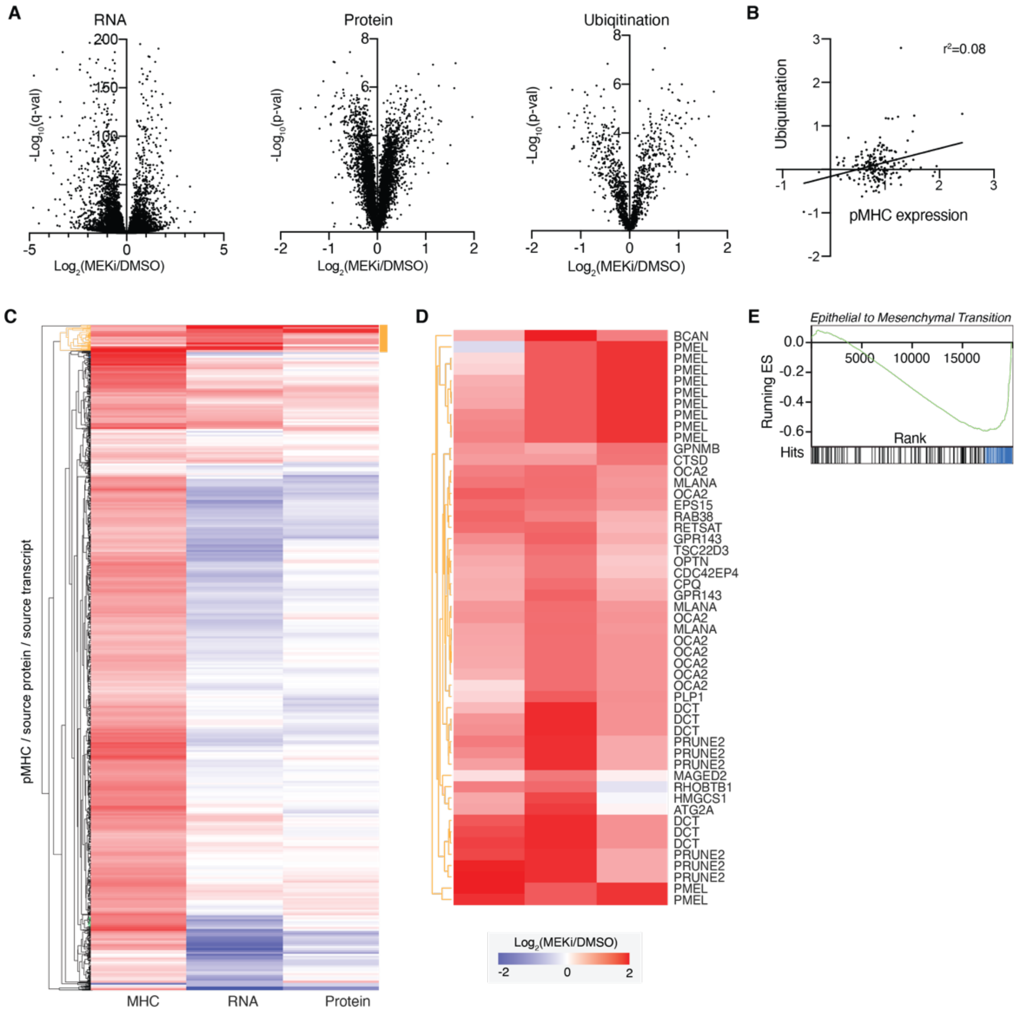


**Fig. S6 Correlation between pMHC, protein, and transcript expression with MEKi treatment.**

**(A)** Volcano plots of changes in RNA and protein expression, and abundance of ubiquitylated peptides. The y-axis represents significance values. Significance: RNA: Wald test, Benjamini Hochberg adjusted. Protein/Ubiquitination: unpaired two-sided T-test. **(B)** Correlation between pMHC expression and ubiquitination levels in SKMEL5 cells +/- 100 nM MEKi. Values represent $Log_2$(MEKi/DMSO). **(C)-(D)**. Hierarchical clustering of pMHC, RNA, and protein expression, represented as the change in expression following 100 nM MEKi. **(D)** displays subcluster highlighted in orange in **(C)**. **(E)** Enrichment plot of EMT genes using for RNA-seq data, SKMEL5 cells +/- 100 nM binimetinib. p & q-values < 0.0001.

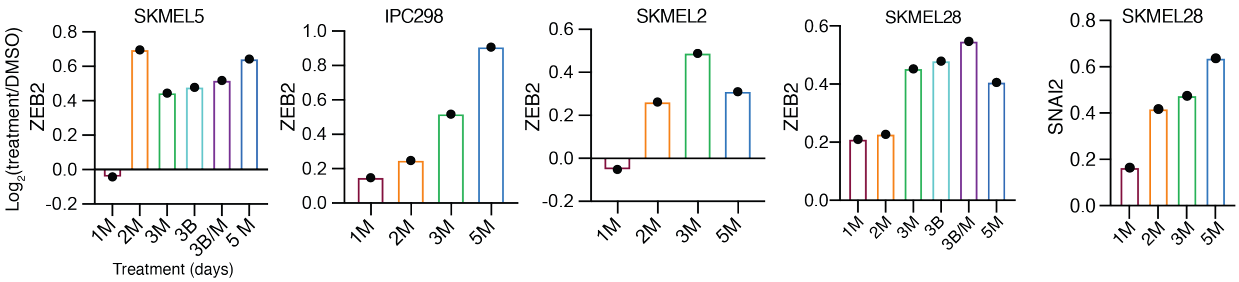


**Fig. S7 pMHC presentation changes in EMT-derived epitopes.**
Maximum change in expression of pMHCs derived from ZEB2 and SNAI2 source proteins from CLX analyses. X-axis describes days of therapy and drug treatment (M=MEKi, binimetinib and B=BRAFi, encorafinib.

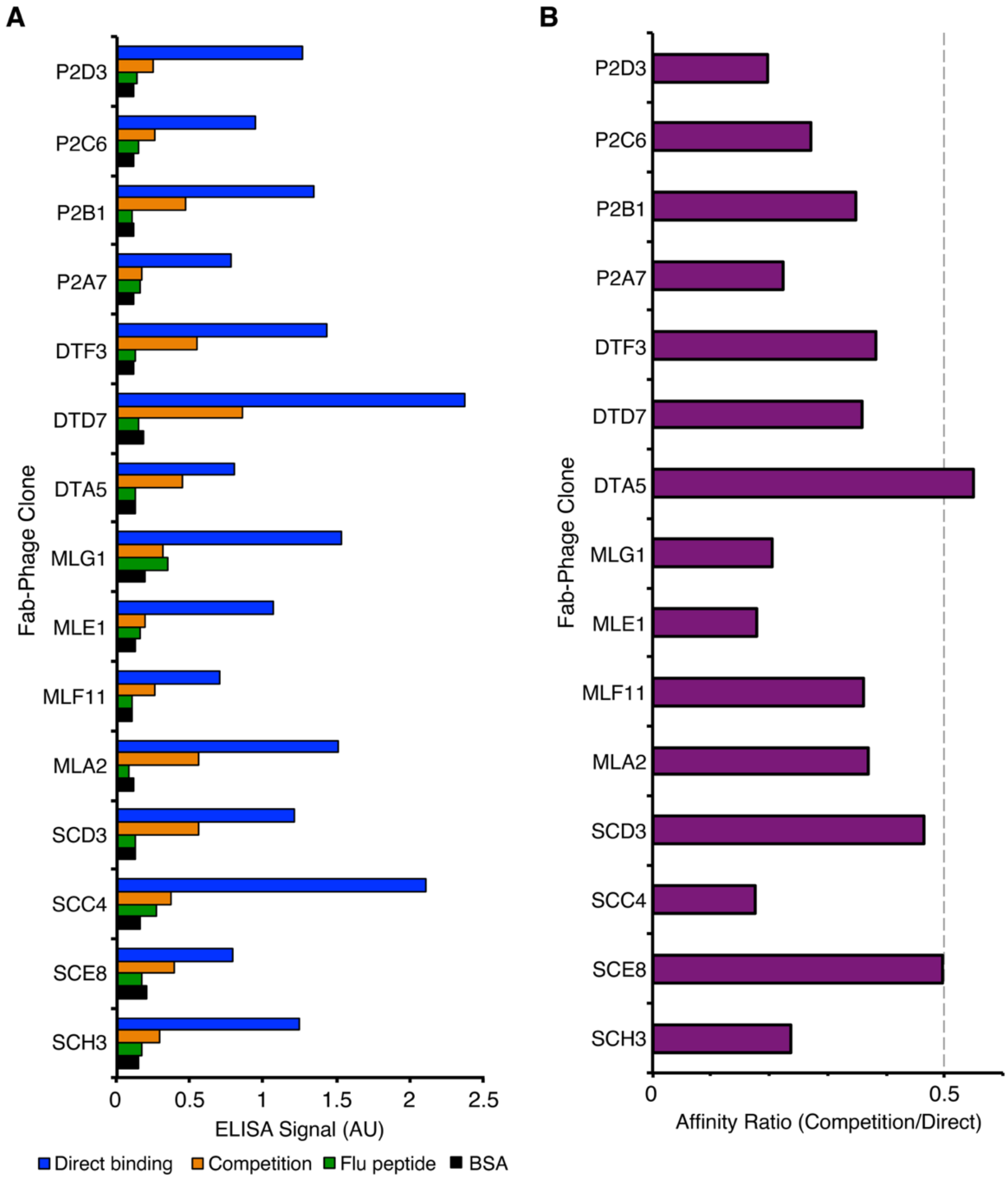


**Fig. S8. Characterization of Fab-phage clones.**
**(A)** Fab-phage ELISA screen of each MHC-peptide complex. Clones with signal above 0.5 AU and competition ratios less than 0.5 were considered high affinity, with predicted relative affinities of <20 nM. Passing clones (green) were further evaluated for binding to the FLU MHC-peptide complex and sequenced. **(B)** ELISA values for all unique Fab-phage clones across all phage-display selections for MHC-peptide complexes.

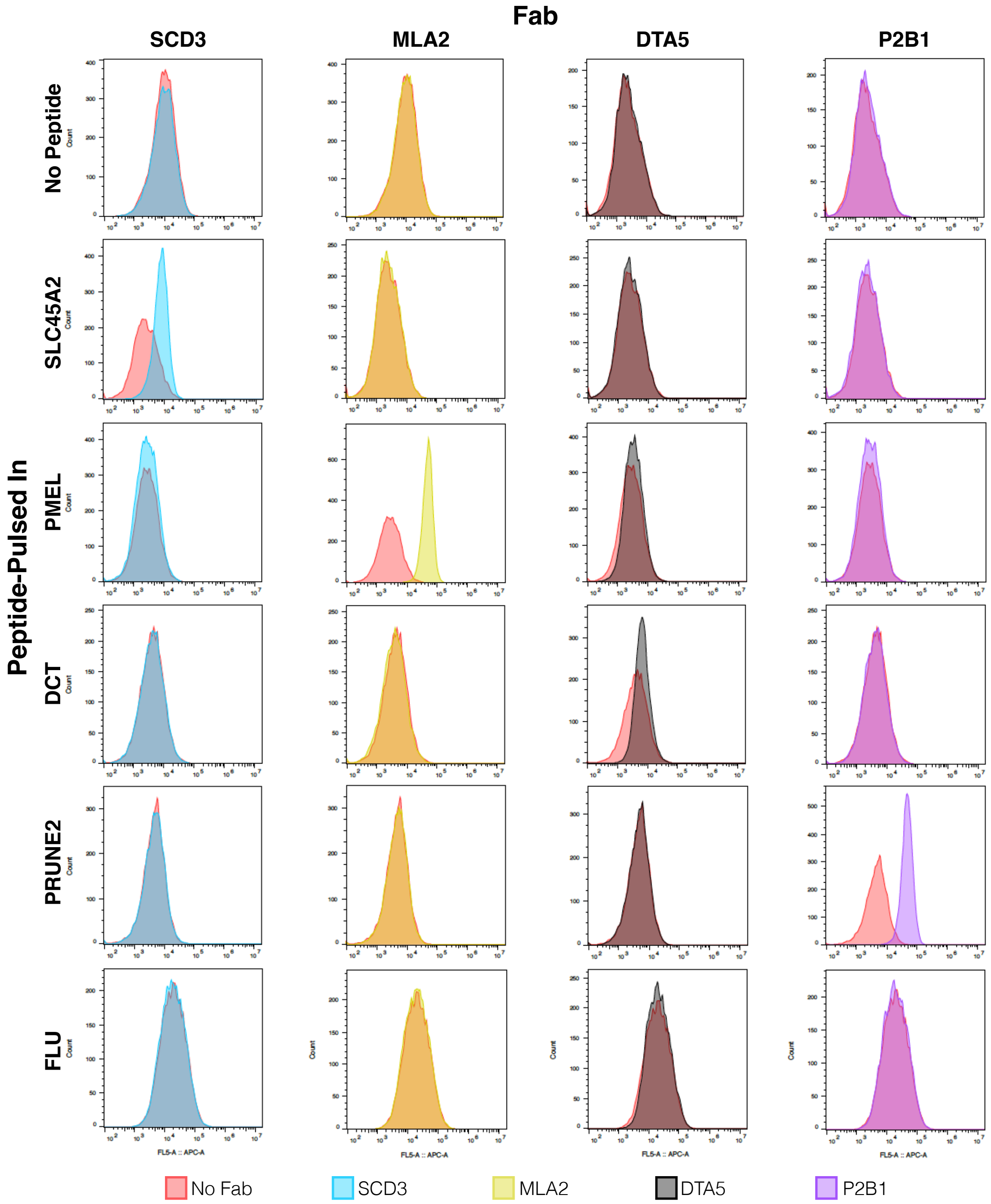


**Fig. S9 Peptide specificity of pMHC-specific Fabs.**
Fluorescence intensity of T2 cells loaded with no peptide (negative control), a decoy FLU peptide, or peptide of interest stained with a pMHC-specific Fab.

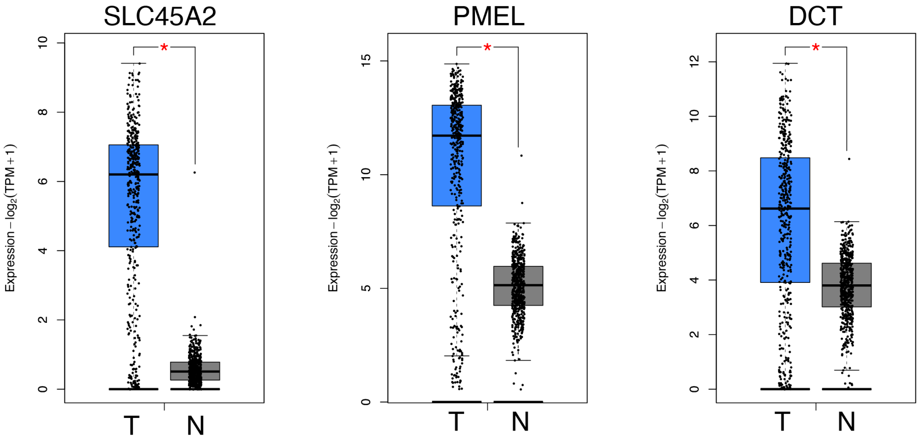


**Fig. S10 Tumor versus normal expression profiles of select TAAs.**
Expression profiles in skin cutaneous melanoma for n=461tumors (T) and n=558 normal tissues (N). *p ≤ 0.01, one way ANOV(A)

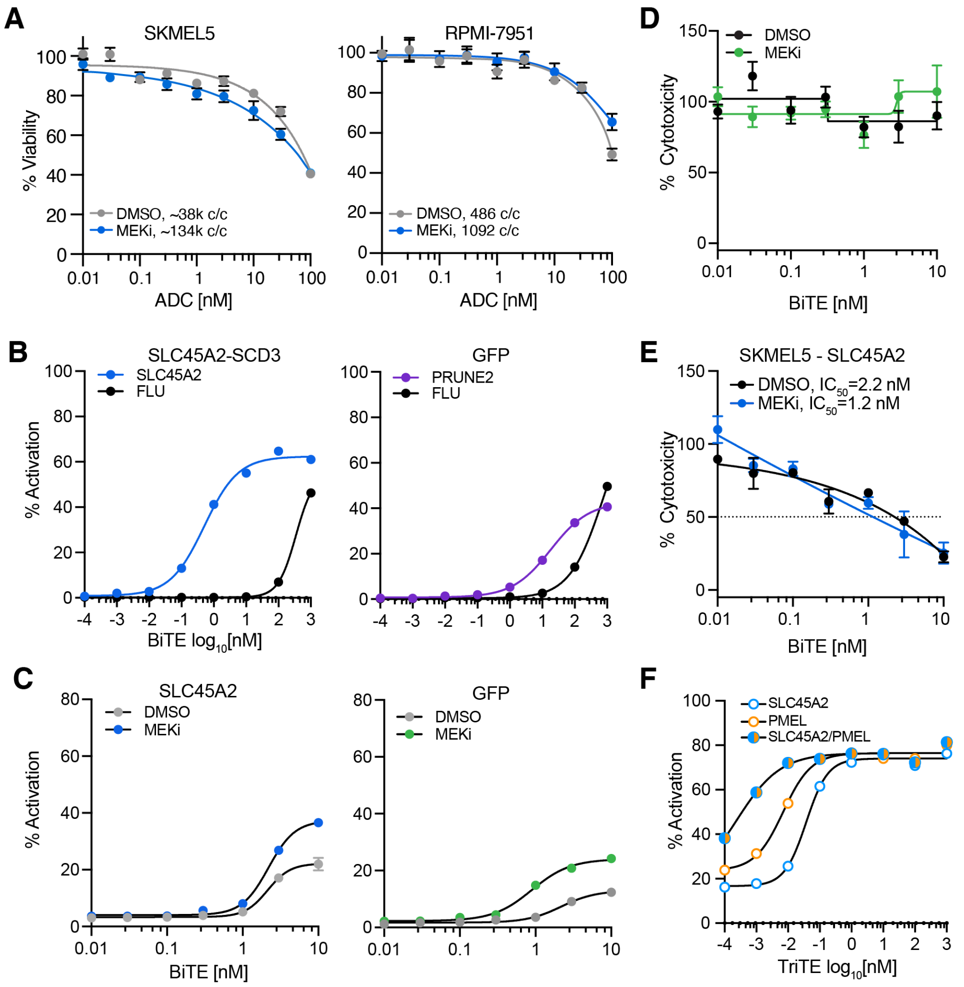


**Fig. S11 Characterization of pMHC-specific ADCs and BiTEs *in vitro*.**
**(A)** Cell viability after 72 hr incubation with SLC45A2-ADC. Error bars represent +/- SD for n=4 biological replicates. Lines represents a four parameter logistic (4PL) nonlinear regression curve. c/c denotes average copies per cell. **(B)-(C)** Percent of GFP+ Jurkat cells following incubation with peptide-pulsed T2 cells **(B)** or SKMEL5 cells **(C)** and a pMHC-specific BiTE or negative control BiTE (anti-GFP) for 24 hours. **(D)** Cell viability (percentage of untreated control) of target cells incubated with normal human T cells (effector:target 2:1) & a negative control anti-GFP BiTE for 48 hours. **(E)** Cell viability (percentage of untreated control) of target cells incubated with normal human T cells (effector:target 2:1) & a pMHC-specific BiTE or 48 hours. **(F)** Percent GFP+ Jurkat cells following incubation with SKMEL5 (DMSO) cells and TriTEs for 24 hours.

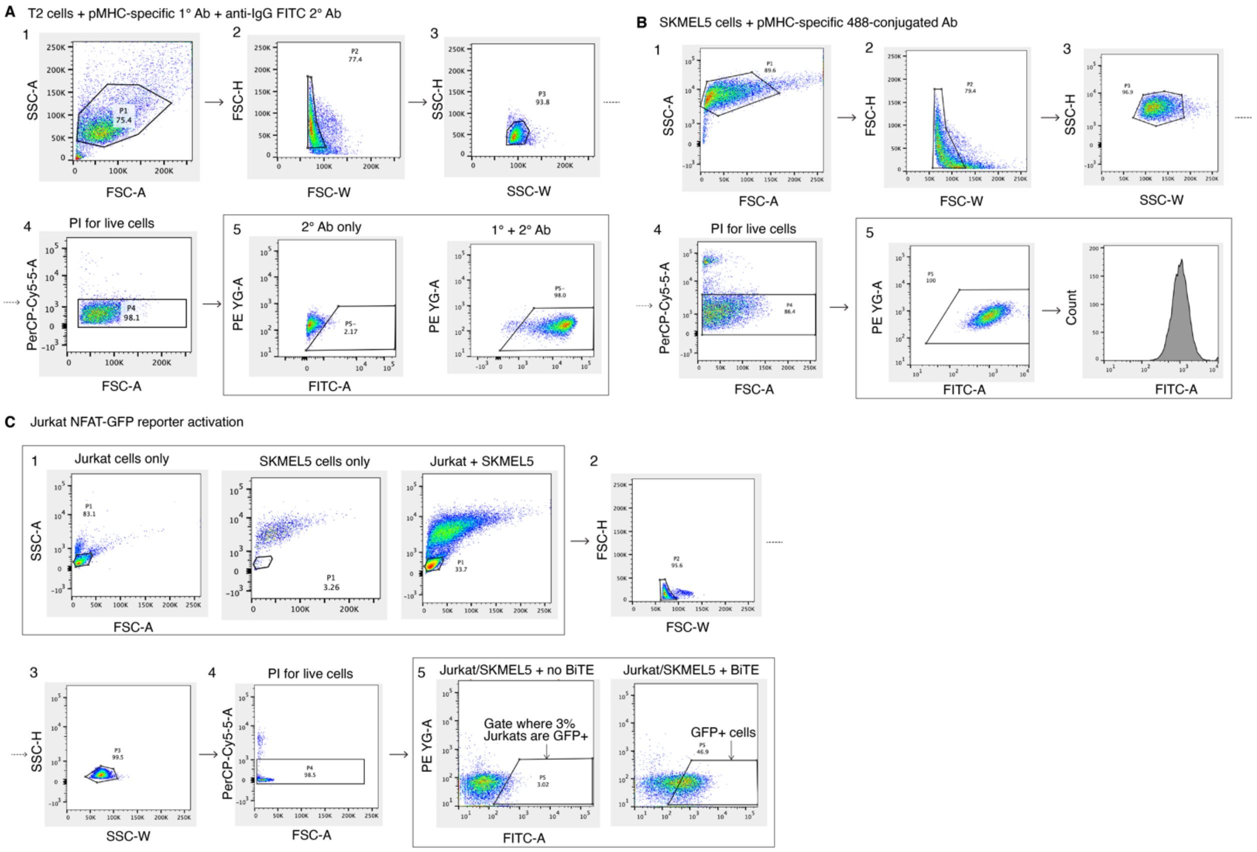


**Figure S12 Flow cytometry gating strategies.**
Flow cytometry gating strategies for **(A)** Peptide-loaded T2 cells + pMHC-specific 1° Ab plus anti-IgG 2° Ab, **(B)** SKMEL5 cells + pMHC-specific Alexa Fluor 488-conjugated Ab, and **(C)** Jurkat NFAT-GFP reporter activation assays.

#### SUPPLEMENTARY METHODS

##### HF-X LC-MS/MS data acquisition

*Chromatography:*

Peptides were resuspended in 0.1% acetic acid and loaded on a precolumn packed in-house (100 µm ID × 10 cm packed with 10 µm C18 beads (YMC gel, ODS-A, 12 nm, S-10 µm, AA12S11)). The precolumn was then washed with 0.1% acetic acid and connected in series to an analytical capillary column with an integrated electrospray tip (~1 µm orifice) with 5µM C18 beads, prepared in house ((50 µm ID × 12 cm with 5 µm C18 beads (YMC gel, ODS-AQ, 12 nm, S-5 µm, AQ12S05)).

*Labeled pMHC analyses:*

Peptides were eluted using a 130-minute gradient with 10-45% buffer B (70% Acetonitrile, 0.2M acetic acid) from 5-100 minutes and 45-55% buffer B from 100-120 minutes at a flow rate of 0.2 mL/min for a flow split of approximately 10,000:1. Peptides were analyzed using a Thermo Q Exactive HF-X Hybrid Quadrupole-Orbitrap mass spectrometer, and data was acquired using Thermo Fisher Scientific Xcalibur version 2.9.0.2923. Standard mass spectrometry parameters were as follows: spray voltage, 2.5 kV; no sheath or auxiliary gas flow; heated capillary temperature, 250 °C.

The HF-X was operated in data-dependent acquisition (DDA) mode for LF and TMT analyses. LF: Full-scan mass spectrometry spectra (mass/charge ratio (*m/z*), 350 to 2,000; resolution, 60,000) were detected in the Orbitrap analyzer after accumulation of ions at $3e^6$ target value with a maximum IT of 50 ms. For every full scan, the top 20 most intense ions were isolated (isolation width of 0.4 *m/z*) and fragmented (collision energy (CE): 28%) by higher energy collisional dissociation (HCD) with a maximum injection time of 350 ms, AGC target $1e^5$, and 30,000 resolution. Charge states < 2 and > 4 were excluded, and dynamic exclusion was set to 45 seconds. TMT: Full-scan mass spectrometry spectra (mass/charge ratio (*m/z*), 400 to 2,000; resolution, 60,000) were detected in the Orbitrap analyzer after accumulation of ions at $3e^6$ target value with a maximum IT of 50 ms. For every full scan, the 20 most intense ions were isolated (isolation width of 0.4 *m/z*) and fragmented (collision energy (CE): 31%) by higher energy collisional dissociation (HCD) with a maximum injection time of 350 ms, AGC target $1e^5$, and 30,000 resolution. Charge states < 2 and > 4 were excluded, and dynamic exclusion was set to 60 seconds.

*Global protein expression profiling:*

Peptides were analyzed using a Thermo Q Exactive HF-X Hybrid Quadrupole-Orbitrap mass spectrometer. Standard mass spectrometry parameters were as follows: spray voltage, 2.5 kV; no sheath or auxiliary gas flow; heated capillary temperature, 250°C. Peptides were eluted with 80% acetonitrile in 0.1% formic acid (buffer B) in following gradient: 0–10% buffer B for 5 min, 10–30% for 100 min, 30–40% for 14 min, 40–60% for 5 min, 60–100% for 2 min, held at 100% of 10 mins, and equilibrated back to 0.1% formic acid. All twenty fractions were analyzed back-to-back.

The HF-X was operated in data-dependent acquisition (DDA) mode. Full-scan mass spectrometry spectra (mass/charge ratio (*m/z*), 300 to 2,000; resolution, 60,000) were detected in the Orbitrap analyzer after accumulation of ions at $3e^6$ target value with a maximum IT of 50 ms. For every full scan, the 15 most intense ions were isolated (isolation width of 0.4 *m/z*) and fragmented (collision energy (CE): 31%) by HCD with a maximum injection time of 350 ms,

AGC target $1e^5$, and 30,000 resolution. Charge states of 1 and >7 were excluded, and dynamic exclusion was set to 20 seconds.

*Ubiquitination analyses:*

The same gradient and standard instrument parameters from global protein expression profiling were used for ubiquitination analyses. The HF-X was operated in data-dependent acquisition (DDA) mode. Full-scan mass spectrometry spectra (mass/charge ratio (m/z), 400 to 1,250; resolution, 60,000) were detected in the Orbitrap analyzer after accumulation of ions at 5e5 target value with a maximum IT of 100 ms. For every full scan, the 20 most intense ions were isolated (isolation width of 0.4 m/z) and fragmented (collision energy (CE): 33%) by HCD with a maximum injection time of 300 ms, AGC target 1e5, and 60,000 resolution. Charge states of <3 and >7 were excluded, and dynamic exclusion was set to 30 seconds. The six fractions were analyzed back-to-back to minimize effects from instrument performance variation.

**Exploris 480 LC-MS/MS data acquisition**

pMHC samples were analyzed using an Orbitrap Exploris 480 mass spectrometer (Thermo Scientific) coupled with an UltiMate 3000 RSLC Nano LC system (Dionex), Nanospray Flex ion source (Thermo Scientific), and column oven heater (Sonation). Samples were resuspended in 0.1% formic acid and directly loaded onto a 10-15 cm analytical capillary chromatography column with an integrated electrospray tip (~1 µm orifice), prepared and packed in house (50 µm ID 1.9 µM C18 beads, ReproSil-Pur). Unless otherwise defined, Standard mass spectrometry parameters were as follows: spray voltage, 2.0 kV; no sheath or auxiliary gas flow; heated capillary temperature, 275 °C.

<u>Labeled DDA pMHC analyses</u>: pMHC elutions were injected in 15-25% fractions for improved coverage of the immunopeptidome. TMT-6/10 chromatography: Peptides were eluted using a gradient with 8-25% buffer B for 50 minutes, 25-35% for 25 minutes, 35-55% for 5 minutes, 55-100% for 2 minutes, hold for 1 minutes, and 100% to 3% for 2 minutes. TMT-Pro chromatography: Peptides were eluted using a gradient with 8-25% buffer B for 50 minutes, 25-45% for 30 minutes, 45-100% for 2 minutes, hold for 1 minutes, and 100% to 3% for 2 minutes.

The Exploris was operated in data dependent acquisition (DDA) mode. Full scan mass spectra (350-1200 m/z, 60,000 resolution) were detected in the orbitrap analyzer after accumulation of $3e^6$ ions (normalized AGC target of 300%) or 25 ms. For every full scan, $MS^2$ were collected during a 3 second cycle time. Ions were isolated (0.4 m/z isolation width) for a maximum of 150 ms or 75% AGC target and fragmented by HCD with 32% CE (TMT-6/10) or 30% (TMT-pro) at a resolution of 45,000. Charge states < 2 and > 4 were excluded, and precursors were excluded from selection for 30 seconds if fragmented n=2 times within 20 second window.

<u>*Isotopologue absolute quantification analyses:*</u>

*Survey analyses of 4H peptides:* Peptides were eluted with 6-25% buffer B for 53 minutes, 25-45% for 12 minutes, 45-97% for 3 minutes, and 97% to 3% for 1 minute. The Exploris was operated in data dependent acquisition (DDA) mode with an inclusion list[2]. Full scan mass spectra (300-1500 m/z, 120,000 resolution) were detected in the orbitrap analyzer after accumulation of $3e^6$ ions or 50 ms. For each full scan, up to 20 ions were subsequently isolated for targets on the inclusion with (+/- 5 ppm of targets m/z) with a minimum intensity threshold of $1e^6$. Ions were collected with a 10s maximum injection time, AGC target: 1000%, and fragmented by HCD with 30% nCE.

*SureQuant-IsoMHC targeted analyses:* Standard mass spectrometry parameters for SureQuant acquisition are as follows: spray voltage: 1.6kV, heated capillary temperature: 280°C. A custom SureQuant acquisition method was built using the Thermo Orbitrap Exploris Series 2.0 software. Full-scan mass spectra were collected with scan range: 350-1200 m/z, AGC target value: $3e^6$, maximum IT: 50 ms, resolution: 120,000. 4H peptides matching the m/z (+/- 3 ppm) and exceeding the defined intensity threshold (1% apex intensity from the survey analysis) were isolated (isolation width 1 m/z) and fragmented by HCD (nCE: 27%) with a scan range: 150-1700 m/z, maximum IT: 10 ms, AGC target: 1000%, resolution: 7,500.

A product ion trigger filter next performs pseudo-spectral matching, where an $MS^2$ scan of the 1H, 2H, 3H, and endogenous peptides are triggered at the defined mass offsets if the 4H trigger peptide contains $n \geq 5$ product ions from the defined list. Scan parameters are the same as the first MS2 scan but with 250 ms max IT, resolution 120,000. The inclusion list, ions for pseudo-spectral matching, and additional method paramters and details have been previously reported.[2]

**LC-MS/MS data analysis:**

All mass spectra were analyzed with Proteome Discoverer (PD, version 2.5) and searched using Mascot (version 2.4) against the human SwissProt database. MS/MS spectra were matched with an initial mass tolerance of 10 ppm on precursor masses and 20 mmu for fragment ions. Data analyses were performed using Matlab version R2019b, and Microsoft Excel version 16.34

*pMHC analyses*: No enzyme was used, static modifications included N-terminal and lysine TMT, and variable modifications included oxidized methionine for all analyses and phosphorylated serine, threonine, and tyrosine for cell treatment analyses. Treatment analyses were also searched against a previously published catalog of over 40,000 predicted antigenic mutations in cancer cell lines.[15] Heavy leucine-containing peptides were searched for separately with heavy leucine (+7), c-terminal amidation, and methionine oxidation as dynamic modifications against a custom database of the synthetic peptide standards. All analyses were filtered with the following criteria: search engine rank =1, isolation interference ≤ 30%, and length between 8 and 15 amino acids. Label-free analyses were filtered with ion score ≥ 20, and labeled samples were filtered with ion score ≥ 15 and percolator q-value ≤ 0.05. Area under the curve (AUC) quantitation was performed using the minora feature detector in PD with match between runs enabled and filtered for ion score ≥ 20.

For TMT-labeled *in vitro* samples, ratios against a reference channel (usually TMT126) were calculated and the median of all ratios for correction hipMHCs was used to determine the final correction parameters. Only PSMs of heavy leucine-coded peptides with an average reporter ion intensity within 10-fold of the interquartile range of endogenous PSM reporter ion intensities were used for correction. To evaluate differences between conditions, the $\log_2$ transformed ratio of arithmetic mean intensity for drug- and DMSO-treated samples (n=3) was calculated. To determine if peptides were significantly increasing, an unpaired, 2-sided t-test was performed, and peptides with p □ 0.05 were considered significantly increasing/decreasing. To evaluate which peptides were significantly enriched above the mean, treated samples were mean centered by dividing the ion intensity of each peptide by the mean fold-change across all peptides, after which a student's 2-tailed t-test was performed on adjusted values. Peptides with a mean-adjusted p-value □ 0.05 were considered significantly enriched. Mean centering was not performed on samples where the mean $\log_2$ fold change was between -0.07 and 0.07.

*Global protein expression profiling:* Enzyme: trypsin, allowing for up to 2 missed cleavages. Cysteine carbamidomethylation, TMT-labeled lysine, and peptide N-termini were searched as fixed modifications, and oxidated methionine was set as a variable modification. PSMs from all fractions were filtered according to search engine rank = 1, ion score ≥ 20, precursor isolation interference ≤30%. Reporter ion abundances for peptides mapping to the same protein were summed, and quantification was corrected by normalizing with the median fold change in TMT abundances over TMT-126 to account for variations in sample input. To determine differences in protein expression, an unpaired, 2-sided t-test was performed, and peptides with $p \leq 0.05$ were considered significantly changing.

*Ubiquitination analysis*: Enzyme: trypsin, allowing for up to 2 missed cleavages. Cysteine carbamidomethylation was set as a static modification, and dynamic modifications were set as diGly-TMT on lysine residues (monoisotopic: 343.20), and N-terminal TMT. PSMs from 6 fractions were filtered to the following criteria: di-Gly modification, search engine rank = 1, ion score ≥ 15, precursor isolation interference ≤ 30%. Reporter ion intensities were summed for PSMs mapping to the same peptide, and the fold change in abundance was calculated by taking the average reporter ion abundance for n=3 replicates per condition. Variation in sample input was account for by normalizing each reporter channel to the median fold change over TMT-126 across all peptides. When multiple peptides mapped to a source protein, the maximal fold change value was used for pMHC comparisons. Significance values were calculated using an unpaired, 2-sided t-test was performed, and peptides with $p \leq 0.05$ were considered significantly changing.

*Isotopologue absolute quantification analyses:*

Peak areas of 6 preselected product ions for each peptide (endogenous and 1-3H isotopologues) were exported from Skyline ((version 20.2.1.28)[16] and summed for all ions quantifiable across the endogenous and isotopologues as previously described.[2] 1-3H peptides were used to generate a calibration curve, from which endogenous pMHC concentrations were determined. Concentrations outside of the standard curve were extrapolated.

#### SUPPLEMENTARY DATA

**Data S1**. In vitro quantitative immunopeptidomics datasets for all cell lines and treatment conditions.

**Data S2**. In vivo Quantitative immunopeptidomics for all cell line xenografts and treatment conditions.

**Data S3.** Transcript expression of SKMEL5 cells +/- 100 nM and 1 uM binimetinib treatment for 72 hours.

**Data S4.** Protein expression of SKMEL5 cells +/- 100 nM binimetinib treatment for 72 hours.

**Data S5**. Peptide ubiquitination levels of SKMEL5 cells +/- 100 nM binimetinib treatment for 72 hours.

**Data S6**. Absolute quantification of 18 tumor associated antigens

#### SUPPLEMENTARY TABLES

**Table S1.** Allelic profile of melanoma cell lines.

| Cell line | HLA-A | HLA-B | HLA-C |
|---|---|---|---|
| SKMEL5 | 02:01, 11:01 | 40:01, 07:02 | 03:04, 07:02 |
| SKMEL28 | 11:01 | 40:01 | 03:04 |
| IPC298 | 24:02, 03:01 | 07:02, 35:02 | 04:01, 07:02 |
| SKMEL2 | 26:01, 03:01 | 38:01, 35:08 | 04:01, 12:03 |
| A375 | 01:01, 02:01 | 44:03, 57:01 | 06:02, 16:01 |
| RPMI-7951 | 01:01, 02:01 | 08:01, 13:02 | 07:01, 06:02 |

**Table S2.** Tumor associated antigen peptide library for enrichment analyses.

| | | | | | |
|---|---|---|---|---|---|
| AAAAAIFVI | ALIHHNTHL | AMLDLLKSV | AVFDGAQVTSK | DLTSFLLSL | ESDPIVAQY |
| AAANIIRTL | ALISKNPV | AMLERQFTV | AVLTKQLLH | DLWKETVFT | ESFSGSLGHL |
| AAFDGRHSQTL | ALKDSVQRA | AMLGTHTMEV | AVMALENNYEV | DNGAKSVVL | ESLFRAVITK |
| AAGIGILTV | ALKDVEERV | AMLGTHTMEVTV | AVQEFGLARFK | DPARYEFLW | ESVMINGKY |
| AARAVFLAL | ALLALTSAV | AMTKDNNLL | AVTNVRTSI | DPKDAEKAI | ETAGPQGPPHY |
| AAVEEGIVLGGG | ALLAVGATK | AMVGAVLTA | AVVDLQGGGHSY | DPSTDYYQEL | ETFTEGQKL |
| ACDGERPTL | ALLEIASCL | AMYDKGPFRSK | AVVGILLVV | DPYKATSAV | ETHLSSKRY |
| ACDPHSGHFV | ALLESSLRQA | ANADLEVKI | AVYGQKEIHRK | DQYPYLKSV | ETILTFHAF |
| AEEAAGIGIL | ALLKDTVYT | ANDPIFVVL | AWISKPPGV | DRASFIKNL | ETLGFLNHY |
| AEEAAGIGILT | ALLMPAGVPL | APAGRPSAS | AWLVAAAEI | DSDPDSFQDY | ETVELQISL |
| AEEHSIATL | ALLNIKVKL | APAGRPSASR | AYACNTSTL | DSFPMEIRQY | ETVSEQSNV |
| AEHIESRTL | ALLPSLSHC | APAGVREVM | AYDFLYNYL | DSFPMEIRQYL | EVAPDAKSF |
| AEINNIIKI | ALLPTALDAL | APDGAKVASL | AYGLDFYIL | DTEFPNFKY | EVAPPASGTR |
| AELESKTNTL | ALMDKSLHV | APLLRWVL | AYIDFEMKI | DVNGLRRVL | EVDPASNTY |
| AELLNIPFLY | ALMEQQHYV | APLQRSQSL | AYTKKAPQL | DVTSAPDNK | EVDPIGHLY |
| AELVHFLLL | ALMVRQARGL | APNTGRANQQM | AYVPQQAWI | DVWSFGILL | EVDPIGHLYIF |
| AEPINIQTW | ALNFPGSQK | APRGVRMAV | CHILLGNYC | DYIGPCKYI | EVDPIGHVY |
| AFLPWHRLF | ALPPPLMLL | APVIKARMM | CIAEQYHTV | DYKSAHKGF | EVFEGREDSVF |
| AFLRHAAL | ALPSFQIPV | AQAEHSITRV | CILGKLFTK | DYLQCVLQI | EVFPLAMNY |
| AGDGTTTATVLA | ALQDIGKNIYTI | AQCQETIRV | CITFQVWDV | DYLQYVLQI | EVFPLAMNYL |
| AGFKGEQGKGEP | ALQKAKQDL | AQDPHSLWV | CLAFPAPAKA | DYLRSVLEDF | EVFQNANFRSF |
| AGYLMELCC | ALRCASPWL | AQPDTAPLPV | CLGHNHKEV | DYPSLSATDI | EVHNLNQLLY |
| AHVDKCLEL | ALREEEEGV | AQYEHDLEVA | CLLSGTYIFA | DYSARWNEI | EVIGRGHFGCVY |
| AIDELKECF | ALSDHHIYL | ARGPESRLL | CLLWSFQTSA | EAAGIGILTV | EVIPYTPAM |
| AIIDPLIYA | ALSEDLLSI | ARGQPGVMG | CLVFLAPAKA | EADPTGHSY | EVIPYTPAMQR |
| AIISGDSPV | ALSVMGVYV | ARHRRSLRL | CLVFPAPAKA | EAFIQPITR | EVIPYTPAMQRY |
| AISANIADI | ALTAVAEEV | ARSVRTRRL | CLVFPAPAKAV | EEFGRAFSF | EVIQWLAKL |
| AIYDHINEGV | ALTDIDLQL | ARTDLEMQI | CMHLLLEAV | EEKLIVVLF | EVISCKLIKR |
| AIYDHVNEGV | ALTEHSLMGM | ASERGRLLY | CMLGDPVPT | EEYLQAFTY | EVISSRGTSM |
| AIYKQSQHM | ALTERSLMGM | ASFDKAKLK | CMTWNQMNL | EEYNSHQSL | EVITSSRTTI |
| AKYLMELTM | ALTPVVVTL | ASGPGGGAPR | CQWGRLWQL | EFKRIVQRI | EVKLSDYKGKYV |
| ALAGLSPV | ALVDAGVPM | ASLDSDPWV | CTACRWKKACQ | EFQKMRRDL | EVLDSLLVQY |
| ALAPAPAEV | ALVSIIKV | ASLIYRRRLMK | CTACRWKKACQR | EGDCAPEEK | EVLLRPGLHFR |
| ALARGAGTVPL | ALWGPDPAAA | ASSTLYLVF | CYMEAVAL | EILGALLSI | EVMSNMETF |
| ALASHLIEA | ALWGPDPAAAF | ASYLDKVRA | CYTWNQMNL | ELAEYLYNI | EVRGDVFPSY |
| ALAVLSVTL | ALWKEPGSNV | ATAGDGLIELRK | DAKNKLEGL | ELAGIGILTV | EVTFVPGLY |
| ALCQNGYHGT | ALWMRLLPL | ATAGIIGVNR | DALVLKTV | ELAPIGHNRMY | EVTSSGRTSI |
| ALCQNGYHGTI | ALWMRLLPLL | ATAQFKINK | DCLVFLAPA | ELFQDLSQL | EVVEKYEIY |
| ALCRWGLLL | ALWPWLLMA | ATATPCWTWLL | DEKQQHIVY | ELHLLQDEEV | EVVHKIIEL |
| ALDEKLLNI | ALWPWLLMA(T) | ATFSSSHRYHK | DELEIKAY | ELHLLQDKEV | EVVRIGHLY |
| ALDGGNKHFL | ALWPWLLMAT | ATGFKQSSK | DEVYQVTVY | ELSDSLGPV | EVYDGREHSA |
| ALDVYNGLL | ALYGDIDAV | ATIIDILTK | DFMIQGGDF | ELTLGEFLK | EYILSLEEL |
| ALEEANADL | ALYLVCGER | ATLPLLCAR | DIKAKMQAS | ELTLGEFLKL | EYLQLVFGI |
| ALENNYEVL | ALYSGVHKK | ATQIPSYKK | DLDVKKMPL | ELVRRILSR | EYLSLSDKI |
| ALFDIESKV | ALYVDSLFFL | ATSPPASVR | DLILELLDL | ELWKNPTAF | EYRGFTQDF |
| ALGDLFQSI | AMAPIKVRL | ATTNILEHY | DLKGFLSYL | EPLARLEL | EYSKECLKEF |
| ALGGHPLLGV | AMAQDPHSL | ATVGIMIGV | DLLSHAFFA | EQYEQILAF | EYSRRHPQL |
| ALIDCNPCTL | AMAQDPHSLWV | AVAANIVLTV | DLPAYVRNL | ERGFFYTPK | EYTAKIALL |
| ALIEVGPDHFC | AMARDPHSL | AVASLLKGR | DLPPPPPLL | ERLERQERL | EYYLQNAFL |
| ALIGGPPV | AMARDPHSLWV | AVCPWTWLR | DLSPGLPAA | ERSPVIQTL | FALQLHDPSGY |

*Table S2 continued*

| | | | | | |
|---|---|---|---|---|---|
| FATPMEAEL | FLPETEPPEM | FVGEFFTDV | GLWRHSPCA | HTMEVTVYHR | IMIHDLCLA |
| FATPMEAELAR | FLPETEPPEML | FVSGSGIAIA | GLYDGMEHL | HTRTPPIIHR | IMIHDLCLV |
| FAWERVRGL | FLPHFQALHV | FVSGSGIATA | GMVTTSTTL | HTYLEPGPVTAQ | IMIHDLCLVFL |
| FEITPPVVL | FLPRNIGNA | FVWLHYYSV | GMWESNANV | HVDSTLLQ | IMLCLIAAV |
| FGLATEKSR | FLQDVMNIL | FYTPKTRRE | GPFGAVNNV | HVDSTLLQV | IMNDMPIYM |
| FGLFPRLCPV | FLRAENETGNM | GADGVGKSA | GPFGPPMPLHV | HVYDGKFLAR | IMPGQEAGL |
| FIASNGVKLV | FLRNFSLML | GADGVGKSAL | GPRESRPPA | HYTNASDGL | IMPKAGLLI |
| FIDKFTPPV | FLRNFSLMV | GAFEHLPSL | GQHLHLETF | IALNFPGSQK | IPSDLERRIL |
| FIDNTDSVV | FLRNLVPRT | GAIAAIMQK | GRAPQVLVL | IARNLTQQL | IPSNPRYGM |
| FIDSYICQV | FLSSANEHL | GASGVGSGL | GSHLVEALY | ICLHHLPFWI | IQATVMIIV |
| FIFPASKVYL | FLSTLTIDGV | GCELKADKDY | GSPATWTTR | IESRTLAIA | IRRGVMLAV |
| FIFSILVLA | FLTGNQLAV | GDFGLATEK | GSSDVIIHR | IEVDGKQVEL | ISGGPRISY |
| FIIENLKAA | FLTKRGGQV | GEISEKAKL | GTAAIQAHY | IIGGGMAFT | ISKPPGVAL |
| FILPVLGAV | FLTKRGRQV | GERGFFYT | GTADVHFER | IIMFDVTSR | ISSVLAGASCPA |
| FINDEIFVEL | FLTKRSGQV | GEVDVEQHTL | GTATLRLVK | IISAVVGIL | ISTQQQATFLL |
| FIQVYEVERA | FLTKRSGQVCA | GFKQSSKAL | GTMDCTHPL | IITEVITRL | ITARPVLW |
| FKNIVTPRT | FLTKRSRQV | GIMAIELAEL | GTMDCTHSL | ILAKFLHWL | ITDFGLAKL |
| FLAELAYDL | FLTPKKLQCV | GIPPAPHGV | GTSSVIVSR | ILAKFLHWLe | ITDFGLARL |
| FLAKLNNTV | FLTPLRNFL | GIPPAPRGV | GTWESNANV | ILAVDGVLSV | ITDQVPFSV |
| FLALIICNA | FLTSGTQFSDA | GIVEQCCTSI | GTYEGLLRR | ILDEKPVII | ITKKVADLVGF |
| FLAPAKAVV | FLWGPRALA | GLAPPQHLIRV | GVALQTMKQ | ILDFGLAKL | ITQPGPLAPL |
| FLAPAKAVVYV | FLWGPRALV | GLASFKSFLK | GVFIQVYEV | ILDKKVEKV | ITQPGPLVPL |
| FLASESLIKQI | FLWGPRAYA | GLCEREDLL | GVLVGVALI | ILDKVLVHL | IVDCLTEMY |
| FLDEFMEGV | FLWSVFMLI | GLEALVPLAV | GVNPVVSYAV | ILDSSEEDK | IVDSLTEMY |
| FLDRFLSCM | FLWSVFWLI | GLEKIEKQL | GVRGRVEEI | ILDTAGREEY | IYMDGTADFSF |
| FLEGNEVGKTY | FLYDDNQRV | GLFDEYLEMV | GVYDGREHTV | ILFGISLREV | KAFLTQLDEL |
| FLFAVGFYL | FLYGALLLA | GLFGDIYLA | GYCASLFAIL | ILGALLSIL | KAFQDVLYV |
| FLFDGSPTY | FLYTLLREV | GLGLPKLYL | GYDQIMPKI | ILHNGAYSL | KAKQDLARL |
| FLFLLFFWL | FMHNRLQYSL | GLGNRWTSRT | GYDQIMPKK | ILIDWLVQV | KALRLSASALF |
| FLGMESCGI | FMNKFIYEI | GLGPVAAV | HAIPHYVTM | ILKDFSILL | KASEKIFYV |
| FLGYLILGV | FMTRKLWDL | GLIEKNIEL | HIAGSLAVV | ILLEAPTGLA | KAYGASKTFGK |
| FLHHLIAEIH | FMTSSWWGA | GLKAGVIAV | HLCGSHLVEA | ILLEAPTGLV | KCDICTDEY |
| FLIIWQNTM | FMTSSWWRA | GLLDKAVSNV | HLFGYSWYK | ILLRDAGLV | KCQEVLAWL |
| FLIVLSVAL | FMTSSWWRAPL | GLLDQVAAL | HLLTSPKPSL | ILLWAARYD | KEADPTGHSY |
| FLLDILGAT | FMVEDETVL | GLLETTVQKV | HLSTAFARV | ILMEHIHKL | KEAGNINTSL |
| FLLENAAYL | FMVELVEGA | GLLGQEGLVEI | HLSYHRLLPL | ILMEHIHKLK | KECVLHDDL |
| FLLENAAYLD | FPALRFVEV | GLLQVHHSCPL | HLSYHWLLPL | ILMEHIHKLKA | KECVLRDDL |
| FLLFIFKVA | FPSDSWCYF | GLMDVQIPT | HLVEALYLV | ILMHCQTTL | KEFEDDIINW |
| FLLGLIFLL | FPYGTTVTY | GLPAGAAAQA | HLWVKNMFL | ILNAMIAKI | KEFEDGIINW |
| FLLKAEVQKL | FQRQGQTAL | GLPGQEGLVEI | HLWVKNVFL | ILPLHGPEA | KEFKRIVQR |
| FLLKLTPLL | FRSGLDSYV | GLPPDVQRV | HLYQGCQVV | ILSAHVATA | KEFTVSGNILTI |
| FLLQMMQICL | FSIDSPDSL | GLPPDVQRVh | HMYHSLYLK | ILSLELMKL | KELEGILLL |
| FLLQMMQVCL | FSYMGPSQRPL | GLPPPPPLL | HPLVFHTNR | ILTVILGVL | KELPSLHVL |
| FLLSLFSLWL | FTHNEYKFYV | GLQHWVPEL | HPRQEQIAL | ILVLASTITI | KEPSEIVEL |
| FLMLVGGSTL | FTWAGKAVL | GLQLGVQAV | HPRYFNQLST | ILYENNVITV | KEWMPVTKL |
| FLMSSWWPNL | FTWAGQAVL | GLREDLLSL | HQILKGGSGTY | ILYENNVIV | KFHRVIKDF |
| FLNQTDETL | FTWEGLYNV | GLREREDLL | HRWCIPWQRL | IMAIELAEL | KFLDALISL |
| FLPATLTMV | FVEHDDESPGL | GLRRVLDEL | HSATGFKQSSK | IMDQVPFSV | KGSGKMKTE |
| FLPEFGISSA | FVEHDLYCTL | GLSPNLNRFL | HSSSHWLRLP | IMFDVTSRV | KIADPICTFI |
| FLPETEPPEI | FVFLRNFSL | GLSTILLYH | HSWITRSEA | IMIGVLVGV | KIDEKTAELK |

*Table S2 continued*

| | | | | | |
|---|---|---|---|---|---|
| KIFDEILVNA | KLQELNYNL | KTWDQVPFS | LLDGTATLRL | LLMPAGVPL | LMLGEFLKL |
| KIFGSLAFL | KLQQKEEQL | KTWDQVPFSV | LLDKAVSNVI | LLMPAGVPLT | LMLQNALTTM |
| KIFSEVTLK | KLQVFLIVL | KTWDQVPFSVSV | LLDRFLATV | LLMWITQCF | LMVLMLAAL |
| KILDAVVAQK | KLSEGDLLA | KTWGQYWQ | LLDTNYNLF | LLNAFTVTV | LMWAKIGPV |
| KINKNPKYK | KLSEQESLL | KTWGQYWQV | LLDTNYNLFY | LLNATIAEV | LNIDLLWSV |
| KIQEILTQV | KLTQINFNM | KVAELVHFL | LLDVAPLSL | LLNLPDKMFL | LNIYEKDDKL |
| KIQRNLRTL | KLVERLGAA | KVAELVRFL | LLDVPTAAV | LLNLPVWVL | LNLPDKMFL |
| KIWEELSVLE | KLVMSQANV | KVFGSLAFV | LLEAPTGLV | LLNQPDKMFL | LPAVVGLSPGEQ |
| KIWEELSVLEV | KLVVVGAVGV | KVHPVIWSL | LLEEMFLTV | LLPENNVLSPV | LPGEVFAI |
| KIYSENLKL | KLYSENLKL | KVIDQQNGL | LLESAFPGGL | LLPPLLEHL | LPHAPGVQM |
| KIYSENLKLA | KLYSENLKLA | KVLEFLAKL | LLFETVMCDT | LLQAEAPRL | LPHNHTDL |
| KIYSENLTL | KLYSENLTL | KVLEHVVRV | LLFGLALIEV | LLQDSVDFSL | LPHSEITTL |
| KIYSENLTLA | KLYSENLTLA | KVLEYVIKV | LLFLLQMMQI | LLQEEEEEL | LPHSSSHWL |
| KLADQYPHL | KMAAFPETL | KVLHELFGMDI | LLFLLQMMQV | LLQEYNWEL | LPLLALLAL |
| KLAEAERVGLHK | KMAELVHFL | KVNIVPVIAK | LLFPYILPPKA | LLQGWVMYV | LPMEVEKNSTL |
| KLAKPLSSL | KMDAEHPEL | KVSAVTLAY | LLFSFAQAV | LLQLGYSGRL | LPPPPPLLDL |
| KLATAQFKI | KMFVKGAPDSV | KVVEFLAML | LLGATCMFV | LLQLYSGRL | LPQKKSNAL |
| KLCKVRKITV | KMFVKGAPESV | KYDCFLHPF | LLGCPVPLGV | LLQMMQICL | LPRWPPPQL |
| KLCPVQLWV | KMISAIPTL | KYIQESQAL | LLGDLFGV | LLQMMQVCL | LPSSADVEF |
| KLDETGNSL | KMLDHEYTT | KYLATASTM | LLGNCLPTV | LLQVHHSCPL | LQSRGYSSL |
| KLDETGNSLK | KMLKSFLKA | KYLKLSSSEL | LLGPGRPYR | LLRGYHQDAY | LRAGRSRRL |
| KLDETGNSLKV | KMNVFDTNL | KYVGIEREM | LLGPTVML | LLSAEVQQHL | LRRYLENGK |
| KLDVGNAEV | KMQASIEKA | LAALPHSCL | LLGRFELIGI | LLSAVLPSV | LSIGTGRAM |
| KLEGLEDAL | KMRRDLEEA | LAAQERRVPR | LLGRNSFEV | LLSDDDVVV | LSRLSNRLL |
| KLFGSLAFV | KMVELVHFL | LALWGPDPAA | LLHVHHSCPL | LLSDEDVAL | LTLGEFLK |
| KLFGVLRLK | KMYAFTLES | LAMPFATPM | LLIADNPQL | LLSDEDVALM | LTLGEFLKL |
| KLGDCIWTYL | KMYAFTLESV | LAPAKAVVYV | LLIDLTSFL | LLSDEDVALMV | LTLTTGEWAV |
| KLGDCIWTYLS | KNKRILMEH | LASEKVYTI | LLIDLTSFLL | LLSDEDVEL | LTYNDFINK |
| KLGDCIWTYPS | KPIVVLHGY | LATEKSRWS | LLIGATIQV | LLSDEDVELM | LTYVSFRNL |
| KLIETYFSK | KPQQKGLRL | LATEKSRWSG | LLIGATIQVT | LLSDEDVELMV | LVALLACLTV |
| KLIGDPNLEFV | KPRQSSPQL | LAVDGVLSV | LLIGATMQV | LLSETVMCDT | LVALLVCLTV |
| KLIKDGLIIRK | KPSGATEPI | LCGSHLVEAL | LLIGATMQVT | LLSGQPASA | LVCGERGFFY |
| KLKHYGPGWV | KPSPPYFGL | LDKVRALEE | LLIGGFAGL | LLSHGAVIEV | LVFGIELMEV |
| KLLDISELDMV | KQDFSVPQL | LEEKKGNYV | LLIKKLPRV | LLSILCIWV | LVFLAPAKAVV |
| KLLEYIEEI | KQDNSTYIMRV | LEEYNSHQSL | LLLDDLLVSI | LLSLFSLWL | LVHFLLLKY |
| KLLGPHVEGL | KQLPEEKQPLL | LEKQLIEL | LLLEAVPAV | LLSPLHCWA | LVLKRCLLH |
| KLLGPHVLGV | KQPAIMPGQSY | LGYGFVNYI | LLLELAGVTHV | LLSPLHCWAV | LVMAPRTVL |
| KLLMVLMLA | KQSSKALQR | LGYGFVNYV | LLLGIGILV | LLSSGAFSA | LVQENYLEY |
| KLLQIQLCA | KRIQEIIEQ | LHHAFVDSIF | LLLGPLGPL | LLTSRLRFI | LVVVGAVGV |
| KLLQIQLCAKV | KRTLKIPAM | LIAHNQVRQV | LLLGTIHAL | LLTTLSNRV | LWMRLLPLL |
| KLLQIQLRA | KSEMNVNMKY | LIFDLGGGT | LLLHCPSKTV | LLVALAIGCV | LYATVIHDI |
| KLLQIQLRAKV | KSLNYSGVK | LIFDLGGGTFD | LLLLDVAPL | LLVSEIDWL | LYAWEPSFL |
| KLLSSGAFSA | KSMNANTITK | LIYDSSLCDL | LLLLTVLTV | LLWWIAVGPV | LYLVCGERGF |
| KLMPPDRTAV | KTCPVQLWV | LIYRRRLMK | LLLPAEVQQHL | LLYKLADLI | LYSACFWWL |
| KLMSPKLYVW | KTIHLTLKV | LKLSGVVRL | LLLPALAGA | LMAGCIQEA | LYSDPADYF |
| KLNVPATFML | KTLGKLWRL | LLAAVAALL | LLLPGPSAA | LMALPPCHAL | LYVDSLFFL |
| KLPNSVLGR | KTLTSVFQK | LLAGIGTVPI | LLLRSPAGV | LMASSPTSI | LYVDSLFFLc |
| KLQAPVQEL | KTPFVSPLL | LLAGPPGV | LLLSAEVQQHL | LMETHLSSK | MALENNYEV |
| KLQATVQEL | KTVDLILEL | LLASSMSSQL | LLMEGVPKSL | LMFWSPSHSCA | MAQKRIHAL |
| KLQEELNKV | KTVNELQNL | LLDDSLVSI | LLMEKEDYHSL | LMGDKSENV | MAVPPCCIGV |

*Table S2 continued*

| | | | | | |
|---|---|---|---|---|---|
| MEGEVWGL | NLFDTAEVYA | PLTSIISAV | RAPPTTPAL | RLLYPDYQI | RTFHHGVRV |
| MEIFIEVFSHF | NLFDTAEVYAA | PPSACSPRF | RASHPIVQK | RLMKQDFSV | RTGEVKWSV |
| MEKEDYHSL | NLFETPVEA | PTLDKVLEL | RAYQQALSR | RLNAALREK | RTIAPIIGR |
| MEVDPIGHLY | NLFLFLFAV | PTLDKVLEV | RCHELTVSL | RLPRIFCSC | RTIPTPLQPL |
| MFPEVKEKG | NLIKLAQKV | QCSGNFMGF | RELEETNQKL | RLQGISPKI | RTKQLYPEW |
| MGNIDSINCK | NLKLKLHSF | QFITSTNTF | REPVTKAEML | RLQREWHTL | RTLAEIAKV |
| MIAVFLPIV | NLLDSLEQYI | QGQHFLQKV | REQFLGALDL | RLQTPMQVGL | RTLDKVLEV |
| MIHDLCLAFPA | NLLEREFGA | QIAKGMSYL | RESEEESVSL | RLQVPVEAV | RTNWPNTGK |
| MIHDLCLVFL | NLLGRFELI | QIEGLKEEL | RFEEKHAYF | RLRAPEVFL | RTTEINFKV |
| MIMQGGFSV | NLLGRFELIGI | QILKGGSGT | RFKMFPEVK | RLRPLCCTA | RVFQGFFTGR |
| MLAVISCAV | NLPDKMFLPGA | QILPLHGPEA | RIAECILGM | RLSCPSPRA | RVHAYIISY |
| MLGDPVPTPT | NLQGSPVYV | QIRPIFSNR | RIDITLSSV | RLSCSSPRA | RVKAPNKSL |
| MLGTHTMEV | NLSALGIFST | QLARQQVHV | RIGQRQETV | RLSSCVPVA | RVLRQEVAAPL |
| MLLAVLYCL | NLSSAEVVV | QLCAKVPLL | RIKDFLRNL | RLTSTNPTM | RVLRQEVEAPL |
| MLLDKNIPI | NLVRDDGSAV | QLCPICRAPV | RILGPGLNK | RLTSTNPTT | RVPGVAPTL |
| MLLKTSEFL | NLWDLTDASVV | QLEERTWLL | RILMEHIHKLK | RLVDDFLLV | RVQEAVESMVK |
| MLLSVPLLLG | NLYPFVKTV | QLFEDNYAL | RINEFSISSF | RLVELAGQSLLK | RVRFFFPSL |
| MLMAQEALAFL | NMQDLVEDL | QLFNHTMFI | RIVQRIKDF | RLWQELSD | RVSLPTSPR |
| MLPSQPTLL | NMVAKVDEV | QLFNKHTMFI | RLAEYQAYI | RLWTTTRPRV | RVTSIRLFEV |
| MLTNSCVKL | NPATPASKL | QLGPTCLSSL | RLARLALVL | RLYDEKQQHI | RVWDLPGVLK |
| MLVGGSTLCV | NPIVVFHGY | QLGPVGGVF | RLASFYDWLP | RLYDEKQQHIVY | RWPSCQKKF |
| MLWSCTFCRI | NPKAFFSVL | QLGRISLLL | RLASSVLRCGK | RLYEMILKR | RYAMTVWYF |
| MLWSCTFCRM | NSELSCQLY | QLIMPGQEA | RLASYLDKV | RLYPWGVVEV | RYCNLEGPPI |
| MLYYPSVSR | NSQPVWLCL | QLLALLPSL | RLDFNLIRV | RMFPNAPYL | RYGSFSVTL |
| MMKMMCIKDL | NTDSPLRY | QLLDGFMITL | RLDQLLRHV | RMLPHAPGV | RYMPPAHRNF |
| MMLPSQPTL | NTYASPRFK | QLLDQVEQI | RLFAFVRFT | RMMEYGTTMV | RYQLDPKFI |
| MMLPSQPTLL | NTYASPRFKf | QLLIKAVNL | RLFFYRKSV | RMMLPSQPTL | RYQQWMERF |
| MMLPSQPTLLT | NVIRDAVTY | QLLKLNVPA | RLFVGSIPK | RMPEAAPPV | SACDVSVRV |
| MMLPSRPTL | NVLHFFNAPL | QLLNSVLTL | RLGGAALPRV | RMTDQEAIQ | SACDVSVRVV |
| MMLPSRPTLL | NVMPVLDQSV | QLMAFNHLV | RLGLQVRKNK | RMTDQEAIQDL | SAFPTTINF |
| MMLPSRPTLLT | NYARTEDFF | QLQGLQHNA | RLGNSLLLK | RNGYRALMDKS | SAGPPSLRK |
| MMQICLHHL | NYKHCFPEI | QLSLLMWIT | RLGPTLMCL | RPHVPESAF | SASVQRADTSL |
| MMQVCLHHL | NYKRCFPVI | QLSSGVSEIRH | RLGPVARTRV | RPKSNIVL | SAWISKPPGV |
| MMSEGGPPGA | NYNNFYRFL | QLVFGIEVV | RLIDLGVGL | RPKSNIVLL | SAYGEPRKL |
| MMYKDILLL | NYSVRYRPGL | QLVIQCEPL | RLIGDAAKNQV | RQAGDFHQV | SEHLDTQKELL |
| MPFATPMEA | PAFSYSFFV | QLYALPCVL | RLLASLQDL | RQFVTQLY | SEIWRDIDF |
| MPFATPMEAEL | PLADLSPFA | QMFFCFKEL | RLLCALTSL | RQKKIRIQL | SEIWRDIDFd |
| MPGEATETV | PLALEGSLQK | QMMQICLHHL | RLLDLAQEGL | RQKRILVNL | SELFRSGLDSY |
| MQLIYDSSL | PLDGGVAAA | QMMQVCLHHL | RLLIKKLPRV | RQLAQEQFFL | SESIKKKVL |
| MSLQRQFLR | PLFDFSWLSL | QQITKTEV | RLLKEYQEL | RQVGDFHQV | SESLKMIF |
| MTSALPIIQK | PLFQVPEPV | QQLDSKFLEQV | RLLPLLALL | RRFFPYYV | SFSYTLLSL |
| MTVDSLVNK | PLHCWAVLL | QRPYGYDQIM | RLLPLLALLAL | RRKWRRWHL | SGMGSTVSK |
| MVIGIPVYV | PLHCWVVLL | QVFPGLLERV | RLLPLWAAL | RRQRRSRRL | SHETVIIEL |
| MVKISGGPR | PLLALLALWG | QVLDLRLPSGV | RLLPLWAALPL | RRRWHRWRL | SHLVEALYLV |
| MVWESGCTV | PLLENVISK | QYSWFVNGTF | RLLQETELV | RSCGLFQKL | SIFDGRVVAK |
| MVYDLYKTL | PLPEAPLSL | RAGLQVRKNK | RLLSDEDVAL | RSDSGQQARY | SIFTWAGKAVL |
| MYIFPVHWQF | PLPPARNGGL | RALAETSYV | RLLSDEDVALM | RSKFRQIV | SIFTWAGQAVL |
| NCLKLLESL | PLPPARNGGLg | RALAKLLPL | RLLSDEDVEL | RSRRVLYPR | SILEDPPSI |
| NLAQDLATV | PLQPEQLQV | RALEEANADLEV | RLLSDEDVELM | RSYHLQIVTK | SIQNYHPFA |
| NLATYMNSI | PLTEYIQPV | RALRLTAFASL | RLLVPTQFV | RSYVPLAHR | SISVLISAL |

*Table S2 continued*

| | | | | | |
|---|---|---|---|---|---|
| SIVKIQSWFRM | SLMSWSAIL | SQGFSHSQM | THFPDETEI | TMTRVLQGV | VLHDDLLE |
| SLAAGVKLL | SLNYSGVKEL | SQKTYQGSY | TIADFWQMV | TPGNRAISL | VLHDDLLEA |
| SLAAYIPRL | SLPGGTAS | SQLTTLSFY | TIHDSIQYV | TPNQRQNVC | VLHELFGMDI |
| SLADEAEVYL | SLPKHSVTI | SQQAQLAAA | TILLGIFFL | TPRLPSSADVEF | VLHWDPETV |
| SLADTNSLAV | SLPPPGTRV | SRASRALRL | TIMIHDLCLA | TPRTPPPQ | VLLESAFPGGL |
| SLADTNSLAVV | SLPRGTSTPK | SRDSRGKPGY | TINPQVSKT | TQPGPLAPL | VLLESAFPGRL |
| SLAMLDLLHV | SLQALKVTV | SRFGGAVVR | TIPTPLQPL | TQPGPLVPL | VLLGMEGSV |
| SLASLLPHV | SLQDVPLAAL | SRFTYTALK | TIRYPDPVI | TRPWSGPYIL | VLLLVLAGV |
| SLAVVSTQL | SLQEEIAFL | SSADVEFCL | TLADFDPRV | TRVLAMAIY | VLLQAGSLHA |
| SLCPWSWRAA | SLQEKVAKA | SSDNYEHWLY | TLAKYLMEL | TSALPIIQK | VLLRHSKNV |
| SLDDYNHLV | SLQKRGIVEQ | SSDYVIPIGTY | TLDEKVAELV | TSDQLGYSY | VLMIKALEL |
| SLDDYNHLVTL | SLQPLALEG | SSFGRGFFK | TLDSQVMSL | TSEHSHFSL | VLNSLASLL |
| SLDKDIVAL | SLQRMVQEL | SSKALQRPV | TLDWLLQTPK | TSEKRPFMCAY | VLNSVASLL |
| SLEEEIRFL | SLQRTVQEL | SSLSLFFRK | TLEEITGYL | TSTTSLELD | VLPDVFIRC |
| SLEENIVIL | SLQSMVQEL | SSPGCQPPA | TLEGFASPL | TTINYTLWR | VLPDVFIRCV |
| SLFEGIDIYT | SLQSTVQEL | SSSGLHPPK | TLGEFLKL | TTLITNLSSV | VLPDVFIRCV |
| SLFEGVDFYT | SLRILYMTL | SSVPGVRLL | TLITDGMRSV | TTNAIDELK | VLQELNVTV |
| SLFGKLQLQL | SLSKILDTV | STALRLTAF | TLKCDCEIL | TVASRLGPV | VLQVGLPAL |
| SLFLGILSV | SLSPLQAEL | STAPPAHGV | TLKKYFIPV | TVFDAKRLIGR | VLQWLPDNRL |
| SLFPNSPKWTSK | SLSRFSWGA | STAPPVHNV | TLLASSMSSQL | TVSGNILTIR | VLQWLSDNRL |
| SLFRAVITK | SLVEELKKV | STDPQHHAY | TLLIGATIQV | TYACFVSNL | VLRDDLLEA |
| SLFVSNHAY | SLWGGDVVL | STIKFQMKK | TLLIGATIQVT | TYLPTNASL | VLRENTSPK |
| SLGEQQYSV | SLWSSSPMA | STLCQVEPV | TLLIGATMQV | TYSEKTTLF | VLRKEEEKL |
| SLGIMAIEL | SLWSSSPMAT | STLQGLTSV | TLLIGATMQVT | VAANIVLTV | VLRQEVAAPL |
| SLGSPVLGL | SLWSSSPMATT | STMPHTSGMNR | TLLLEGVMAA | VAELVHFLL | VLSVNVPDV |
| SLGWLFLLL | SLYHVYEVNL | STPPPGATRV | TLLNLPDKMFL | VAVKAPGFGD | VLTSESMHV |
| SLIAAAAFCLA | SLYKFSPFPL | STSQEIHSATK | TLLNQPDKMFL | VCGERGFFYT | VLVEGSTRI |
| SLIKQIPRI | SLYQLENYC | STVASRLGPV | TLLPATMNI | VCLHHLPFWI | VLVPPLPSL |
| SLKLLESLTPI | SLYSFPEPEA | STVASWLGPV | TLLSNIQGV | VEETPGWPTTL | VLWDRTFSL |
| SLLDRFLATV | SMCRFSPLTL | SVAQQLLNGK | TLMSAMTNL | VEGSGELFRW | VLYGPDAPTV |
| SLLGLALLAV | SMLIRNNFL | SVASLLPHV | TLPGYPPHV | VEIEERGVKL | VLYPRVVRR |
| SLLGQLSGQV | SMPPPGTRV | SVASTITGV | TLPPAWQPFL | VFLPCDSWNL | VLYRYGSFSV |
| SLLKFLAKV | SMPQGTFPV | SVGSVLLTV | TLPPRPDHI | VIMPCSWWV | VLYRYGSFSVTL |
| SLLLELEEV | SMSKEAVAI | SVHSLHIWSL | TLRTGEVKWSV | VISNDVCAQV | VMALENNYEV |
| SLLMWITQA | SMSSQLGRISL | SVKPASSSF | TLSSRVCCRT | VIVMLTPLV | VMIIVSSLAV |
| SLLMWITQC | SNDGPTLI | SVPQLPHSSSHW | TLTNIAMRPGL | VIWEVLNAV | VMLDKQKEL |
| SLLMWITQCFL | SPASSRTDL | SVQGIIIYR | TLTTGEWAV | VLAGGFFLL | VMNILLQYV |
| SLLNLPVWV | SPAVDKAQAEL | SVSPVVHVR | TLWVDPYEV | VLAGVGFFI | VMNILLQYVV |
| SLLNLPVWVLM | SPGSGFWSF | SVVKIQSWFRM | TLYEAVREV | VLASIEAEL | VPGWGIALL |
| SLLPAIVEL | SPHPVTALL | SVYDFFVWL | TLYNPERTITV | VLASIEAELPM | VPLDCVLYRY |
| SLLQHLIGL | SPLFQRSSL | SYLDKVRA | TMASTSVSRSA | VLASIEPEL | VPRSAATTL |
| SLLQSREYSSL | SPQNLRNTL | SYLDSGIHF | TMESMNGGKLY | VLASIEPELPM | VPYGSFKHV |
| SLLQSRGYSSL | SPRFSPITI | SYRNEIAYL | TMGGYCGYL | VLCSIDWFM | VRIGHLYIL |
| SLLSGDWVL | SPRPPLGSSL | SYTRLFLIL | TMHSLTIQM | VLDFRLPSGV | VRLGSLSTK |
| SLLSLPVWV | SPRWWPTCL | TALRLTAFASL | TMKIYSENLTL | VLDGLDVLL | VRSRRCLRL |
| SLLSLPVWVLM | SPSKAFASL | TCQPTCRSL | TMKLYSENLTL | VLEGMEVV | VSDFGGRSL |
| SLLSPLHCWA | SPSSNRIRNT | TEAASRYNL | TMLARLASA | VLFGLGFAI | VSLLSLPVWV |
| SLLSPLHCWAV | SPSVDKARAEL | TETEAIHVF | TMLGRRAPI | VLFSSDFRI | VTLLIGATIQV |
| SLLTSSKGQLQK | SPTSSRTSSL | TFDYLRSVL | TMLGRRPPI | VLFYLGQY | VTLLIGATMQV |
| SLMASSPTSI | SQFGGGSQY | TFPDLESEF | TMNGSKSPV | VLFYLGQYI | VTTDIQVKV |

*Table S2 continued*

| | | |
|---|---|---|
| VVEGTAYGL | YLDPAQQNL | YVDPVITSI |
| VVHFFKNIV | YLEPGPVTA | YVFTLLVSL |
| VVLGVVFGI | YLEYRQVPV | YVIPIGTYGQM |
| VVMSWAPPV | YLFSEEITSG | YYNAFHWAI |
| VVMVNQGLTK | YLGSYGFRL | YYSVRDTLL |
| VVPCEPPEV | YLIELIDRV | YYWPRPRRY |
| VVPEDYWGV | YLLKPVQRI | RVASPTSGVK |
| VVQNFAKEFV | YLLLRVLNI | |
| VVTGVLVYL | YLLPAIVHI | |
| VVVGAVGVG | YLLPEAEEI | |
| VYDFFVWLHY | YLLQGMIAAV | |
| VYDYNCHVDL | YLMDTSGKV | |
| VYFFLPDHL | YLNDHLEPWI | |
| VYGIRLEHF | YLNKEIEEA | |
| VYLDKFIRL | YLQGMIAAV | |
| VYSDADIFL | YLQLVFGIEV | |
| VYVKGLLAKI | YLQQNTHTL | |
| WATLPLLCAR | YLSGANLNL | |
| WGPDPAAA | YLSGANLNLG | |
| WLDEVKQAL | YLSGANLNV | |
| WLDPNETNEI | YLVGNVCIL | |
| WLEYYNLER | YLVPQQGFFC | |
| WLLPLWAAL | YLYDRLLRI | |
| WLLPLWAALPL | YLYGQTTTY | |
| WLPFGFILI | YMDGTMSQV | |
| WLPKILGEV | YMFDVTSRV | |
| WLQYFPNPV | YMFPNAPYL | |
| WLSLKTLLSL | YMIAHITGL | |
| WLSLLFKKL | YMIDPSGVSY | |
| WMRLLPLLAL | YMIMVKCWMI | |
| WQYFFPVIF | YMIPSIRNSI | |
| WYEGLDHAL | YMMPVNSEV | |
| WYQTKYEEL | YMNGTMSQV | |
| YAVDRAITH | YMNSIRLYA | |
| YEDIHGTLHL | YPFKPPKV | |
| YEGSPIKVTL | YQGSYGFRL | |
| YGHSGQASGLY | YQLDPKFIV | |
| YGYDNVKEY | YRPRPRRY | |
| YIDEQFERY | YRYGSFSVTL | |
| YIFAVLLVCV | YSDHQPSGPYY | |
| YIGEVLVSV | YSLEYFQFV | |
| YLAMPFATPME | YSLKLIKRL | |
| YLAPENGYL | YSWMDISCWI | |
| YLCDKVIPG | YTCPLCRAPV | |
| YLCDKVVPG | YTDFHCQYV | |
| YLCSGSSYF | YTDFVGEGL | |
| YLCSGSSYFV | YTDQPSTSQIAY | |
| YLDLFGDPSV | YTLDRDSLYV | |
| YLDLLFQIL | YTMKEVLFY | |
| YLDLLFQILL | YVDFREYEYY | |

**Table S3**. Custom library of tumor associated antigen source proteins.

| | | | | | |
|---|---|---|---|---|---|
| ABCA1 | BING4 | CLTC | DUSP22 | GNTK | IMP3 |
| ABCA6 | BIRC5 | CLYBL | EEF2 | GPC3 | INPP5D |
| ABCC3 | BIRC7 | CML28 | EFTUD2 | GPCPD1 | INS |
| ABCD3 | BIRC8 | CNMD | EGFR | GPNMB | INSM2 |
| ABL1 | BIRC9 | COA1 | EHD2 | GPR143 | INTS11 |
| ACPP | BRAF | COL2A1 | EIF2S3 | HAO2 | INTS13 |
| ACRBP | BST2 | COL4A3 | EIF3D | HAUS3 | IQGAP2 |
| ACTB | BTBD2 | COL6A2 | ELAC2 | HBD | IRS2 |
| ACTN4 | BTG1 | CORO1A | ELAVL1 | HCG | ITGAL |
| ADAM17 | BTK | COX2 | ELAVL4 | HDAC1 | ITGAM |
| ADAMTSL5 | C18orf21 | CPSF | EML6 | HDGF | ITGB2 |
| ADRP | C2CD4A | CPSF1 | ENAH | HEPACAM | ITGB8 |
| AFP | C5 | CPVL | EPCAM | HHAT | KAAG1 |
| AIM2 | CA9 | CR2 | EPHA2 | HIFPH3 | KCNAB1 |
| AIMP1 | CACNG | CRABP1 | ERAP1 | HIST4H4 | KDM2B |
| AKAP13 | CADH3 | CRNKL1 | ERBB2 | HIVEP1 | KDM5B |
| ALDH1A1 | CALCA | CSAG2 | ERVK3-1 | HLA-A | KDM5C |
| ALDOA | CALR | CSF1 | ETV5 | HLA-B | KDM5D |
| ALK | CAMP | CSF3R | EVI2B | HLA-DOB | KDR |
| ALYREF | CASP5 | CSNK1A1 | EZH2 | HLA-DPA1 | KIAA1551 |
| AML1 | CASP8 | CSPG4 | EZR | HMMR | KIF20A |
| AMZ2 | CCDC110 | CT83 | FAM136A | HMOX1 | KLK10 |
| ANKRD30A | CCL3 | CTAG1A | FASN | HMSD | KLK3 |
| ANO7 | CCL3L1 | CTDP1 | FBXW11 | HNF4G | KLK4 |
| ANXA1 | CCLA2 | CTNNB1 | FCER1A | HNRNPL | KRAS |
| APOBEC3H | CCNA1 | CTPS1 | FDPS | HNRNPLL | KRI1 |
| ARF1 | CCNB1 | CTSH | FGF5 | HNRNPR | KRT16 |
| ARHGAP15 | CCND1 | CYP1B1 | FGF6 | HOXD3 | KRT18 |
| ARHGAP25 | CCNI | CYP21A2 | FLT3 | HPN | KRT6C |
| ARHGAP4 | CD19 | CYP2A6 | FLT3LG | HPSE | KTN1 |
| ARHGAP45 | CD274 | CYP2A7 | FMOD | HSDL1 | LAGE1 |
| ARL4D | CD33 | CYP2C8 | FMR1NB | HSP90AB1 | LAGE3 |
| ART4 | CD48 | CYP2C9 | FNDC3B | HSPA1A | LAS1L |
| ART5 | CD69 | CYP2D6 | FOLH1 | HSPA1B | LCK |
| ASH1L | CD79B | CYPB | FOXO1 | HSPA1L | LCP2 |
| ATIC | CDC5L | DAPK2 | G3BP1 | HSPA6 | LGALS1 |
| ATP2A3 | CDCA7L | DCT | G6PC2 | HSPB1 | LGALS3BP |
| ATXN10 | CDH13 | DDX21 | GAD2 | HSPD1 | LGSN |
| B2A2 | CDK12 | DDX3Y | GAGE1 | ICAM3 | LPGAT1 |
| B3A2 | CDK4 | DDX5 | GAS7 | ICE | LRMP |
| BA46 | CDKN1A | DKK1 | GATA2 | IDNK | LRRC8A |
| BAD | CDKN2A | DLAT | GC | IDO1 | LTB |
| BAGE1 | CDR2 | DMD | GCGR | IER3 | LY6K |
| BCAP31 | CEACAM5 | DMXL1 | GEMIN4 | IFG2BP3 | LYN |
| BCHE | CELF6 | DNAJC2 | GFAP | IFI30 | MAG |
| BCL2 | CELSR1 | DNMBP | GINS1 | IFI6 | MAGEA1 |
| BCL2A1 | CENPM | DNMT1 | GLRX3 | IGF2BP2 | MAGEA12 |
| BCL2L4 | CEP55 | DOCK2 | GLS | IGF2BP3 | MAGEB2 |
| BFAR | CLCA2 | DOK2 | GNAO1 | IL13RA2 | MAGEC1 |
| BID | CLP | DSE | GNL3L | IL2RG | MAGEC2 |

*Table S3 continued*

| | | | | |
|---|---|---|---|---|
| MAGEC2 | NELFA | PSMB1 | SIRT2 | UBE2C |
| MAGED2 | NFYC | PSMB10 | SLC25A5 | UBE2D2 |
| MAGEE1 | NISCH | PTHLH | SLC30A8 | UGT2B17 |
| MAGEF1 | NLRP5 | PTPN11 | SLC41A3 | UQCR10 |
| MAGEF1 | NOB1 | PTPN21 | SLC45A2 | UQCRH |
| MALL | NONO | PTPRC | SLC45A3 | USP11 |
| MAP4K1 | NPM1 | PTPRN | SLCO2A1 | USP9X |
| MARK3 | NQO1 | PTTG1IP | SNRNP70 | USP9Y |
| MATN2 | NRAS | PUM3 | SNRPD1 | UTY |
| MBP | NUDCD1 | PWWP3A | SNX14 | VEGFA |
| MC1R | NUF2 | PXDN | SOX10 | VENTXP1 |
| MCF2 | NUF3 | RAB38 | SP110 | VGF |
| MCM5 | NUP210 | RAN | SPA17 | VIM |
| MCMBP | NUP37 | RASGRF1 | SPARC | VIPR1 |
| MDK | OCA2 | RASGRP2 | SPATA5L1 | VPS13B |
| MDM2 | OGT | RASSF10 | SSX1 | VSIG10L |
| MDN1 | OS9 | RBAF600 | SSX2 | WNK2 |
| ME1 | P2RX5 | RBBP4 | STAT1 | WT1 |
| MED23 | PAK2 | RBL2 | STEAP1 | XAGE1B |
| MED24 | PARP10 | RFA1 | SUGT1 | XBP1 |
| MET | PARP3 | RGS5 | SUPT5H | ZFAND5 |
| METTL21A | PASD1 | RHOC | SYNGR1 | ZFHX3 |
| MFGE8 | PAX3 | RINT1 | TAG1 | ZFP36L1 |
| MICA | PAX5 | RNF19B | TALDO1 | ZFY |
| MLANA | PCDH11Y | RNF43 | TBC1D22A | ZMYM4 |
| MMP2 | PCDH20 | RPA1 | TCHH | ZNF395 |
| MMP7 | PDGFRA | RPL10A | TEK | |
| MOK | PFKM | RPL19 | TEP1 | |
| MPL | PGK1 | RPS2 | TERT | |
| MRPL19 | PHB | RPS4Y1 | TG | |
| MS4A1 | PHRF1 | RPSA | TGRBR2 | |
| MSCP | PIM1 | RUBCNL | THEM6 | |
| MSLN | PLAC1 | SAGE1 | TMCO1 | |
| MT-ATP6 | PLIN2 | SART1 | TMED4 | |
| MT-CO2 | PLP1 | SART3 | TMSB10 | |
| MTRR | PMEL | SASH1 | TMSB4Y | |
| MUC1 | POP1 | SCGB2A2 | TOP1 | |
| MUC16 | PP2A | SCGB2A7 | TOP2A | |
| MUC5AC | PPFIBP1 | SCRN1 | TP53 | |
| MUM2 | PPIB | SEC31A | TP53I11 | |
| MUM3 | PPP1R3B | SELL | TPBG | |
| MYH1 | PRAME | SELPLG | TPO | |
| MYH2 | PRDM1 | SEPT2 | TRIM22 | |
| MYH9 | PRDX2 | SEPT6 | TRIM68 | |
| MYO1G | PRDX5 | SERPINB5 | TTK | |
| N4BP2 | PRELID1 | SF1 | TTN | |
| N4BP2L1 | PRKCB | SFMBT1 | TYMS | |
| NACA2 | PRTN3 | SGT1B | TYR | |
| NCF4 | PSD4 | SH3GLB2 | TYRP1 | |
| NECTIN4 | PSMA3 | SIRPD | UBD | |

**Table S4.** CLX treatment groups and dosing schedule. Groups 6 and 7 were not included in SKMEL2/IPC298 studies (*NRAS* mutant).

| **Study groups** | **Schedule** | **Route** |
|---|---|---|
| 1. Vehicle (1%CMC/0.5%Tween80) | BID1-3, 2hr | PO |
| 2. 3.5 mg/kg MEK162 | QD, 2hr | PO |
| 3. 3.5 mg/kg MEK162 | BID1-2, 2hr | PO |
| 4. 3.5mg/kg MEK162 | BID1-3, 2hr | PO |
| 5. 3.5mg/kg MEK162 | BID1-5, 2hr | PO |
| 6. 20mg/kg LGX818 | QD1-3, 2hr | PO |
| 7. 3.5 mg/kg MEK162/ 20mg/kg LGX818 | BID1-3, 2hr / QD1-3 | PO |
